# Nominally non-responsive frontal and sensory cortical cells encode task-relevant variables via ensemble consensus-building

**DOI:** 10.1101/347617

**Authors:** Michele N. Insanally, Ioana Carcea, Rachel E. Field, Chris C. Rodgers, Brian DePasquale, Kanaka Rajan, Michael R. DeWeese, Badr F. Albanna, Robert C. Froemke

## Abstract

Neurons recorded in behaving animals often do not discernibly respond to sensory input and are not overtly task-modulated. These nominally non-responsive neurons are difficult to interpret and are typically neglected from analysis, confounding attempts to connect neural activity to perception and behavior. Here we describe a trial-by-trial, spike-timing-based algorithm to reveal the hidden coding capacities of these neurons in auditory and frontal cortex of behaving rats. Responsive and nominally non-responsive cells contained significant information about sensory stimuli and behavioral decisions, and network modeling indicated that nominally non-responsive cells are important for task performance. Sensory input was more accurately represented in frontal cortex than auditory cortex, via ensembles of nominally non-responsive cells coordinating the behavioral meaning of spike timings on correct but not error trials. This unbiased approach allows the contribution of all recorded neurons - particularly those without obvious task-modulation - to be assessed for behavioral relevance on single trials.

---

Spike trains recorded from the cerebral cortex of behaving animals can be complex, highly variable from trial-to-trial, and therefore challenging to interpret. A fraction of recorded cells typically exhibit trial-averaged responses with obvious task-related features and can be considered ‘classically responsive’, such as neurons with tonal frequency tuning in the auditory cortex or orientation tuning in the visual cortex. Another population of responsive cells are modulated by multiple task parameters (‘mixed selectivity cells’), and have recently been shown to have computational advantages necessary for flexible behavior^1^·. However, a substantial number of cells have variable responses that fail to demonstrate any obvious trial-averaged relationship to task parameters^2–5^. These ‘nominally non-responsive’ neurons are especially prevalent in frontal cortical regions but can also be found throughout the brain, including primary sensory cortex^4–6^. These response categories are not fixed but can be dynamic, with some cells apparently becoming nominally non-responsive during task engagement without impairing behavioral performance^7–9^. The potential contribution of these cells to behavior remains to a large extent unknown and represents a major conceptual challenge to the field^2^.

How do these nominally non-responsive cells relate to behavioral task variables on single trials? While there are sophisticated approaches for dissecting the precise correlations between responsive cells and task structure^3,4,10–12^ there is still a need for complementary and straightforward analytical tools for understanding any and all activity patterns encountered^1,3,4^. Moreover, most behavioral tasks produce dynamic activity patterns throughout multiple neural circuits, but we lack unified methods to compare activity across different regions, and to determine to what extent these neurons might individually or collectively perform task-relevant computations. To address these limitations, we devised a novel trial-to-trial analysis using Bayesian inference that evaluates the extent to which single-unit responses and ensembles encode behavioral task variables.

## Results

### Nominally non-responsive cells prevalent in auditory and frontal cortex during behavior

We trained 15 rats on an audiomotor frequency recognition go/no-go task^8,13–15^ that required them to nose poke to a single target tone for food reward and withhold from responding to other non-target tones (**Fig. 1a**). Tones were 100 msec in duration presented sequentially once every 5-8 seconds at 70 dB sound pressure level (SPL); the target tone was 4 kHz and non-target tones ranged from 0.5-32 kHz separated by one octave intervals. After a few weeks of training, rats had high hit rates to target tones and low false alarm rates to non-targets, leading to high d’ values (mean performance shown in **Fig. 1b**; each individual rat included in this study shown in **Supplementary Fig. 1**).

**Figure 1.**
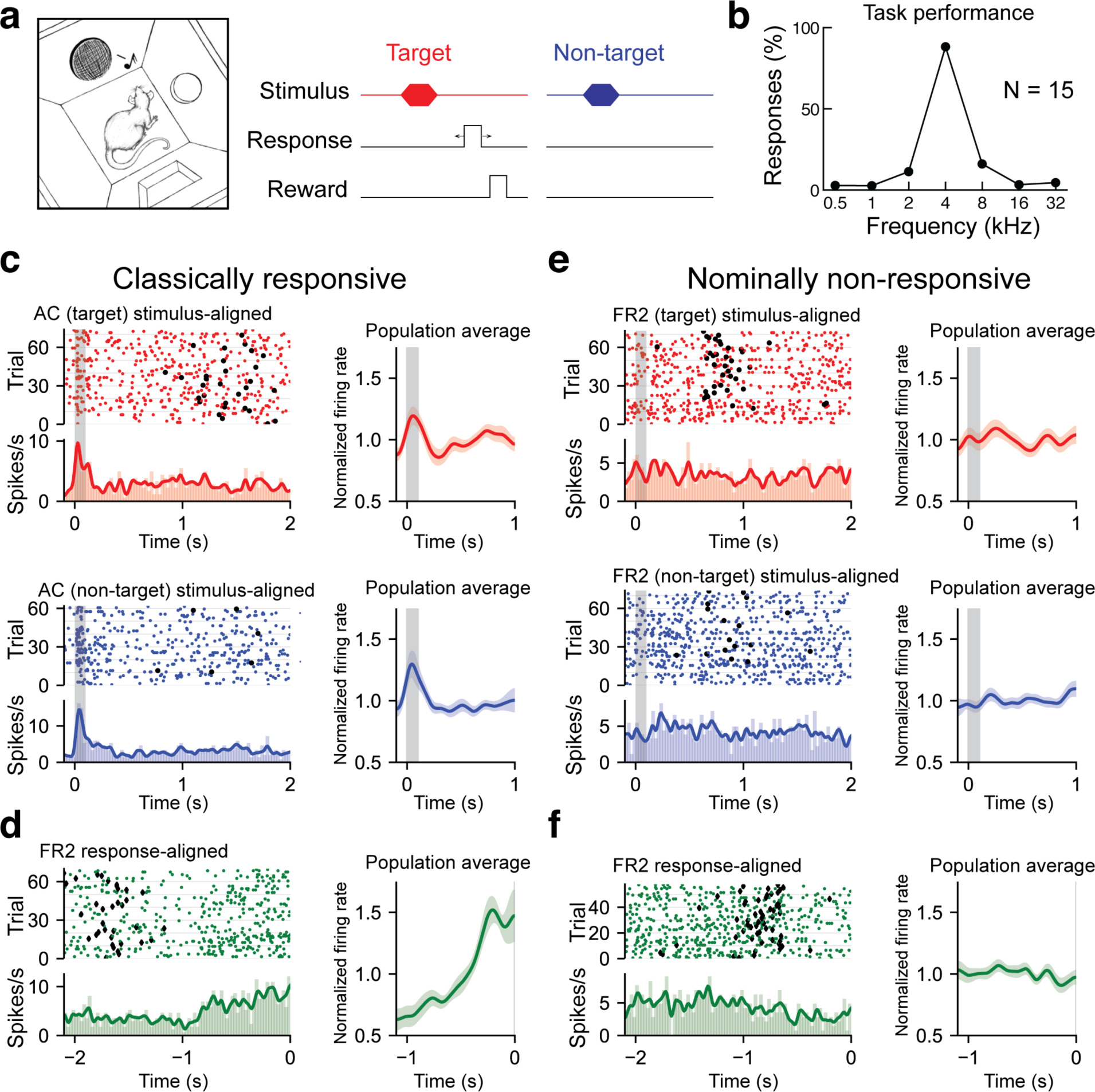
Recording from AC or FR2 during go/no-go audiomotor task. **a**. Behavioral schematic for the go/no-go frequency recognition task. Animals were rewarded with food for entering the nose port within 2.5 seconds after presentation of a target tone (4 kHz) or given a 7-second time-out if they incorrectly responded to non-target tones (0.5, 1, 2, 8,16, or 32 kHz). **b**. Behavioral responses (nose pokes) to target and non-target tones (hit rates: 88 ± 7%, false alarms: 7 ± 5%, N=15 rats). **c**. Left, AC unit with significant tone modulated responses during target trials (red; top panel, average evoked spikes = 0.55) and non-target trials (blue; bottom panel, average evoked spikes = 0.92). Rasters of individual trials as well as the firing rate histogram and moving average are shown. Histograms of average firing rate during a trial were constructed using 25 ms time bins. A moving average of the firing rate was constructed using a Gaussian kernel with a 20 ms standard deviation. Black circles represent behavioral responses. Right, population averages for all target (n=23) or nontarget (n=34) responsive singe-units from AC. **d**. Left, FR2 unit with ramping activity (green; ramp index = 2.82). Trials here are aligned to response time. Diamonds indicate stimulus onset. Right, population average for all ramping single-units from FR2 (n=21). **e**. Left, FR2 unit that was not significantly modulated during target trials (red; average evoked spikes = .041, p<.001, 2,000 bootstraps). Black circles here represent behavioral responses. Right, population averages for all target (n=44) or non-target (n=44) non-responsive single-units from FR2 **f**. Left, FR2 unit lacking ramping activity (green, ramp index = −1.0, p<.001, 2,000 bootstraps). Right, population average for all non-ramping single-units from FR2 (n=44).

To correctly perform this task, animals must first recognize the stimulus and then execute an appropriate motor response. We hypothesized that two brain regions important for this behavior are the auditory cortex (AC) and frontal cortical area 2 (FR2). Many but not all auditory cortical neurons respond to pure tones with reliable, short-latency phasic responses^6,16–21^. These neurons can process sound in a dynamic and context-sensitive manner, and AC cells are also modulated by expectation, attention, and reward structure, strongly suggesting that AC responses are important for auditory perception and cognition^4,21–24^. Previously we found that the go/no-go tone recognition task used here is sensitive to AC neuromodulation and plasticity^13^. In contrast, FR2 is not thought to be part of the canonical central auditory pathway, but is connected to many other cortical regions including AC^25,26^. This region has recently been shown to be involved in orienting responses, categorization of perceptual stimuli, and in suppressing AC responses during movement^10,25,27^. These characteristics suggest that FR2 may be important for goal-oriented behavior.

We first asked if activity in AC or FR2 is required for animals to successfully perform this audiomotor task. We implanted cannulas into AC or FR2 (**Supplementary Fig. 2**), and infused the GABA agonist muscimol bilaterally into AC or FR2, to inactivate either region prior to testing behavioral performance. We found that task performance was impaired if either of these regions was inactivated, although general motor functions, including motivation or ability to feed were not impaired (**Supplementary Fig. 3**; for AC p=0.03; for FR2 p=0.009 Student’s paired two-tailed t-test). Thus activity in both AC and FR2 may be important, perhaps in different ways, for successful task performance.

Once animals reached behavioral criteria (hit rates ≥70% and d’ values ≥1.5), they were implanted with tetrode arrays in either AC or FR2 (**Supplementary Fig. 4**). After recovery, we made single-unit recordings from individual neurons or small ensembles of 2-8 cells during task performance. The trial-averaged responses of some cells exhibited obvious task-related features: neuronal activity was tone-modulated compared to inter-trial baseline activity (**Fig. 1c**) or gradually increased over the course of the trial as measured by a ramping index (**Fig. 1d**; hereafter referred to as ‘ramping activity’). However, 60% of recorded cells were nominally non-responsive in that they were neither tone modulated nor ramping according to statistical criteria (**Fig. 1e,f; Supplementary Fig. 5**; 64/103 AC cells and 43/74 FR2 cells from 15 animals had neither significant tone-modulated activity or ramping activity; pre and post-stimulus mean activity compared via bootstrapping and considered significant when p<0.05; ramping activity measured with linear regression and considered significant via bootstrapping when p<0.05 and r>0.5; for overall population statistics see **Supplementary Fig. 6**).

### Novel single-trial, ISI-based algorithm for decoding non-responsive activity

Given that the majority of our recordings were from nominally non-responsive cells, we developed a general method for interpreting neural responses even when trial-averaged responses were not obviously task-modulated which allowed us to compare coding schemes across different brain regions (here, AC and FR2). The algorithm is agnostic to the putative function of neurons as well as the task variable of interest (here, stimulus category or behavioral choice).

Our algorithm uses the interspike intervals (ISIs) of individual neurons to decode the stimulus category (target or non-target) or behavioral choice (go or no-go) on each trial. In principle, any response property could be used with our method; however, we chose the ISI because its distribution could vary between task conditions even without changes in the firing rate building on previous work demonstrating that the ISI distribution contains complementary information to the firing rate^28–30^. The distinction between the ISI distribution and trial-averaged firing rate is subtle, yet important. While the ISI is obviously closely related to the instantaneous firing rate, decoding with the ISI distribution is not simply a proxy for using the time-varying, trial-averaged rate. To demonstrate this we constructed three model cells: a stimulus-evoked cell with distinct target and non-target ISI distributions (**Fig. 2a**), a stimulus-evoked cell with identical ISI distributions (**Fig. 2b**), and a nominally non-responsive cell with distinct target and non-target ISI distributions (**Fig. 2c**). These models clearly demonstrate that trial-averaged rate modulation can occur with or without corresponding differences in the ISI distributions and cells without apparent trial-averaged rate-modulation can nevertheless have distinct ISI distributions. Taken together, these examples demonstrate that the ISI distribution and trial-averaged firing rate capture different spike train statistics. This has important implications for decoding non-responsive cells that by definition do not exhibit large firing rate modulations but nevertheless may contain information hidden in their ISI distributions.

**Figure 2.**
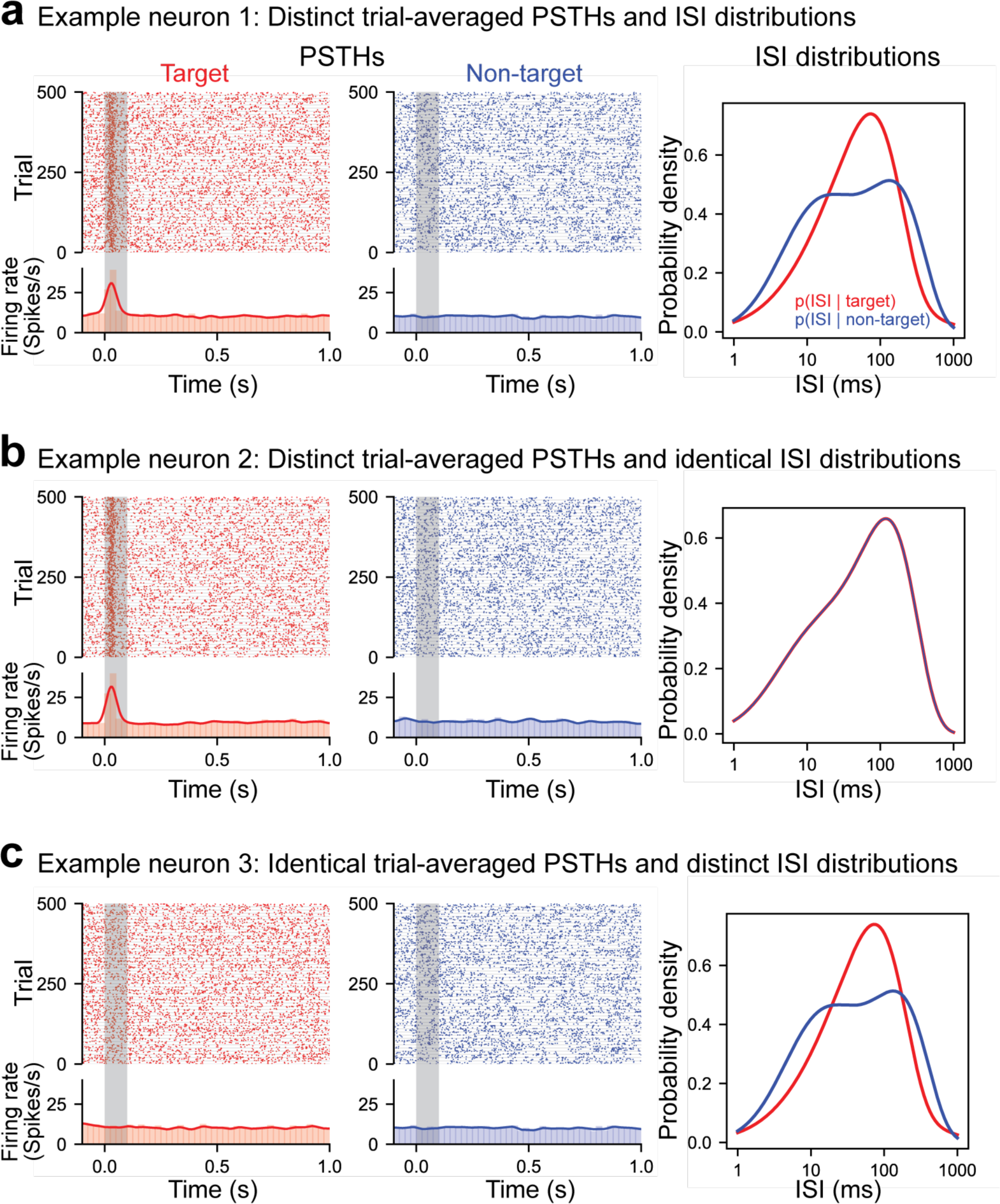
ISIs capture information distinct from trial-averaged rate. Three simulated example neurons demonstrating that differences in the ISI are not necessary for differences in the trial-averaged firing rate to occur (and vice versa). Each trial was generated by randomly sampling from the appropriate conditional ISI distribution. Evoked responses were generated by shifting trials without altering the ISI distributions such that one spike during stimulus presentation is found at approximately 30 ms (with a variance of 10 ms). **a**. Example neuron with both an evoked target response and a difference in the conditional ISI distributions on target and non-target trials. **b**. Example neuron with an evoked target response but identical conditional ISI distributions. **c**. Example nominally non-responsive neuron with no distinct trial-averaged activity relative to the pre-stimulus period that nevertheless is generated by distinct ISI distributions.

For each recorded neuron, we built a library of ISIs observed during target trials and a library for non-target trials from a set of ‘training trials’. Two different cells from AC are shown in **Fig. 3** and **Supplementary Fig. 7a-d**, and another cell from FR2 is shown in **Supplementary Fig. 7e-h**. These libraries were used to infer the probability of observing an ISI during a particular trial type (**Fig. 3b,c; Supplementary Fig. 7c,g**; left panels show target in red and non-target in blue). These conditional probabilities were inferred using non-parametric statistical methods to minimize assumptions about the underlying process generating the ISI distribution and better capture the heterogeneity of the observed ISI distributions (**Fig. 3b; Supplementary Fig.7c,g**). We verified that our observed distributions were better modeled by non-parametric methods rather than standard parametric methods (e.g. rate-modulated Poisson process; **Supplementary Fig. 8**). Specifically, we found the distributions using Kernel Density Estimation where the kernel bandwidth for each distribution was set using 10-fold cross-validation. We then used these training set probability functions to decode a spike train from a previously unexamined individual trial from the set of remaining ‘test trials’. This process was repeated 124 times using 10-fold cross-validation with randomly generated folds.

**Figure 3.**
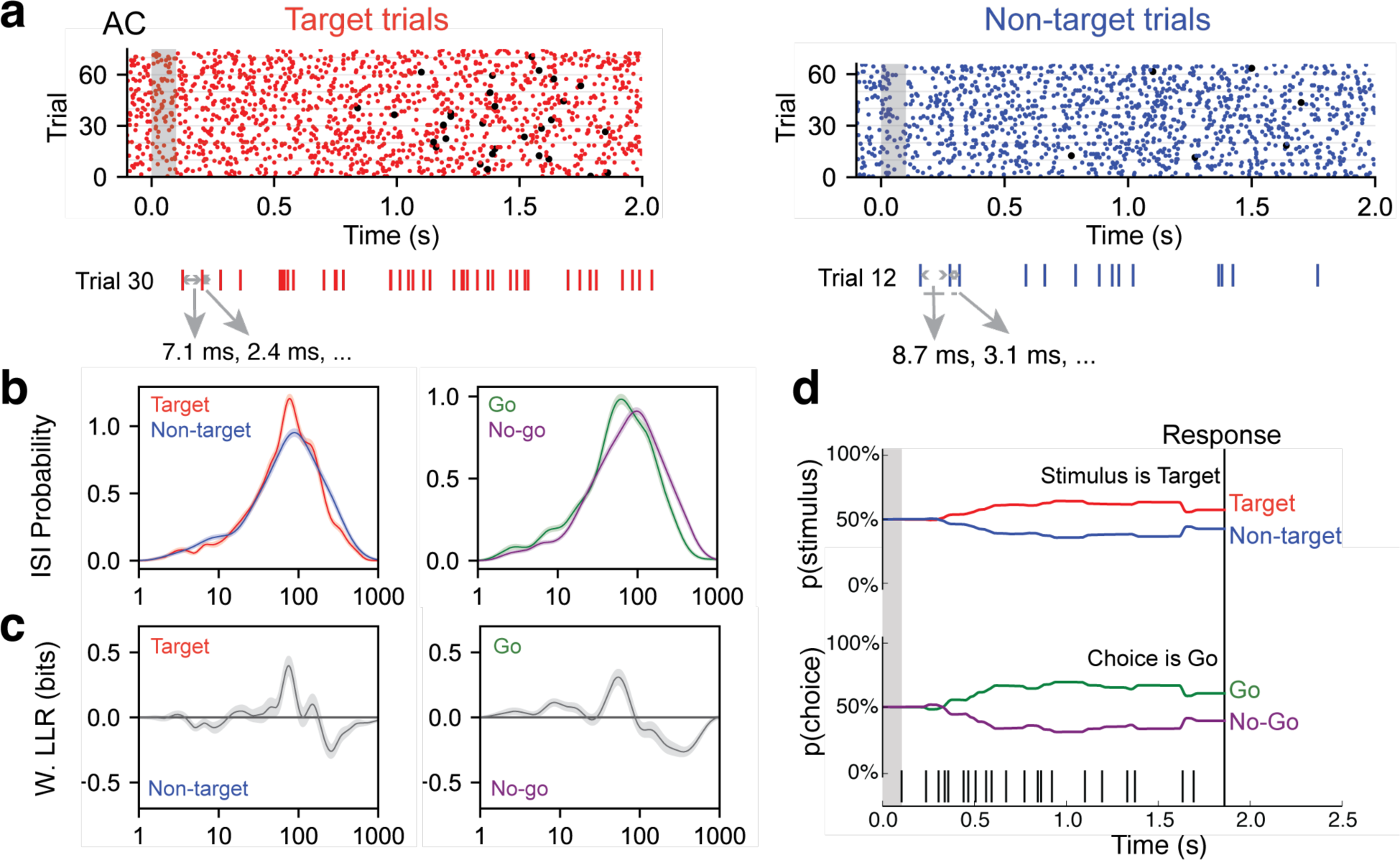
ISI-based algorithm for decoding behavioral variables from AC and FR2 single-units. **a**. Single-unit activity was first sorted by task condition, here for target trials (red) and non-target trials (blue). All ISIs following stimulus onset and before behavioral choice were aggregated into libraries for each condition (average response time is used on no-go trials) as shown for a sample trial. **b**. Probability of observing a given ISI on each condition was generated via Kernel Density Estimation on libraries from **a**. Left, target (red) and non-target (blue) probabilities. Right, go (green) and no-go (purple). **c**. Relative differences between the two stimulus conditions (or choice conditions) was used to infer the actual stimulus category (or choice) from an observed spike train, in terms of weighted log likelihood ratio (W. LLR) for stimulus category (p(ISI)*(log2p(ISI|target) - log2(ISI|non-target)); on left) and behavioral choice (p(ISI)*(log p2(ISI|go) - log2(ISI|no-go)); on right). When curve is above zero the ISI suggests target (go) and when below zero the ISI suggests non-target (no-go). **d**. Probability functions from c were used as the likelihood function to estimate the prediction of a spike train on an individual trial (bottom). Bayes’ rule was used to update the probability of a stimulus (top) or choice (bottom) as the trial progressed and more ISIs were observed. The prediction for the trial was assessed at the end of the trial.

Importantly, while the probabilities of observing particular ISIs on target and non-target trials were similar (**Fig. 3b; Supplementary Fig. 7c,g**), small differences between the curves carried sufficient information to allow for decoding. To characterize these differences, we used the weighted log likelihood ratio (W. LLR; **Fig. 3c; Supplementary Fig. 7c,g**) to clearly represent which ISIs suggested target (W. LLR >0) or non-target (W. LLR <0) stimulus categories. Our algorithm relies only on statistical differences between task conditions; therefore, the W. LLR summarizes all spike timing information necessary for decoding. Similar ISI libraries were also computed for behavioral choice categories (**Fig. 3b,c; Supplementary Fig. 7c,g**; right panels show go decision in green and no-go in purple). These examples clearly illustrate that the relationship between the ISIs and task variables can be non-monotonic: in the cell shown in **Fig. 3**, short ISIs (ISI <50 msec) indicated non-target, medium ISIs (50 msec < ISI < 100 msec) indicated target, and longer ISIs indicated non-target (100 msec < ISI). This non-monotonic relationship would be impossible to capture with a simple ISI or firing rate threshold.

The algorithm uses the statistical prevalence of certain ISI values under particular task conditions (in this case the ISIs accompanying stimulus category or behavioral choice), to infer the task condition for each trial. Each trial begins with equally uncertain probabilities about the stimulus categories (i.e., p(target) = p(non-target) = 50%). As each ISI is observed sequentially within the trial, the algorithm applies Bayes’ rule to update p(target|ISI) and p(non-target|ISI) using the likelihood of the ISI under each stimulus category (p(ISI|target) and p(ISI|non-target) (**Fig. 3b-d**). As shown for one trial of the example cell in **Fig. 3d**, ISIs observed between 0-1.0 seconds consistently suggested the presence of the target tone, whereas ISIs observed between 1.0-1.4 seconds suggested the non-target category thereby also necessarily reducing the belief that a target tone was played (**Fig. 3d**, top trace). After this process was completed for all ISIs in the particular trial, we obtained the probability of a non-target tone and a target tone as a function of time during the trial (**Fig. 3d**). The prediction for the entire trial p(target|ISI) is evaluated at the end of the trial (in the example trial, p(target|ISI) = 61%; **Fig 3d**). This process is repeated for the behavioral choice (**Fig 3b-d**; right panels; trials separated according to go, no-go; probabilities of ISIs in each condition generated; conditional probabilities used as likelihood function to predict behavioral choice on a given trial). The single-trial decoding performance of each neuron is then averaged over all trials as a measure of the overall ability of each neuron to distinguish behavioral conditions (**Fig. 4a**).

**Figure 4.**
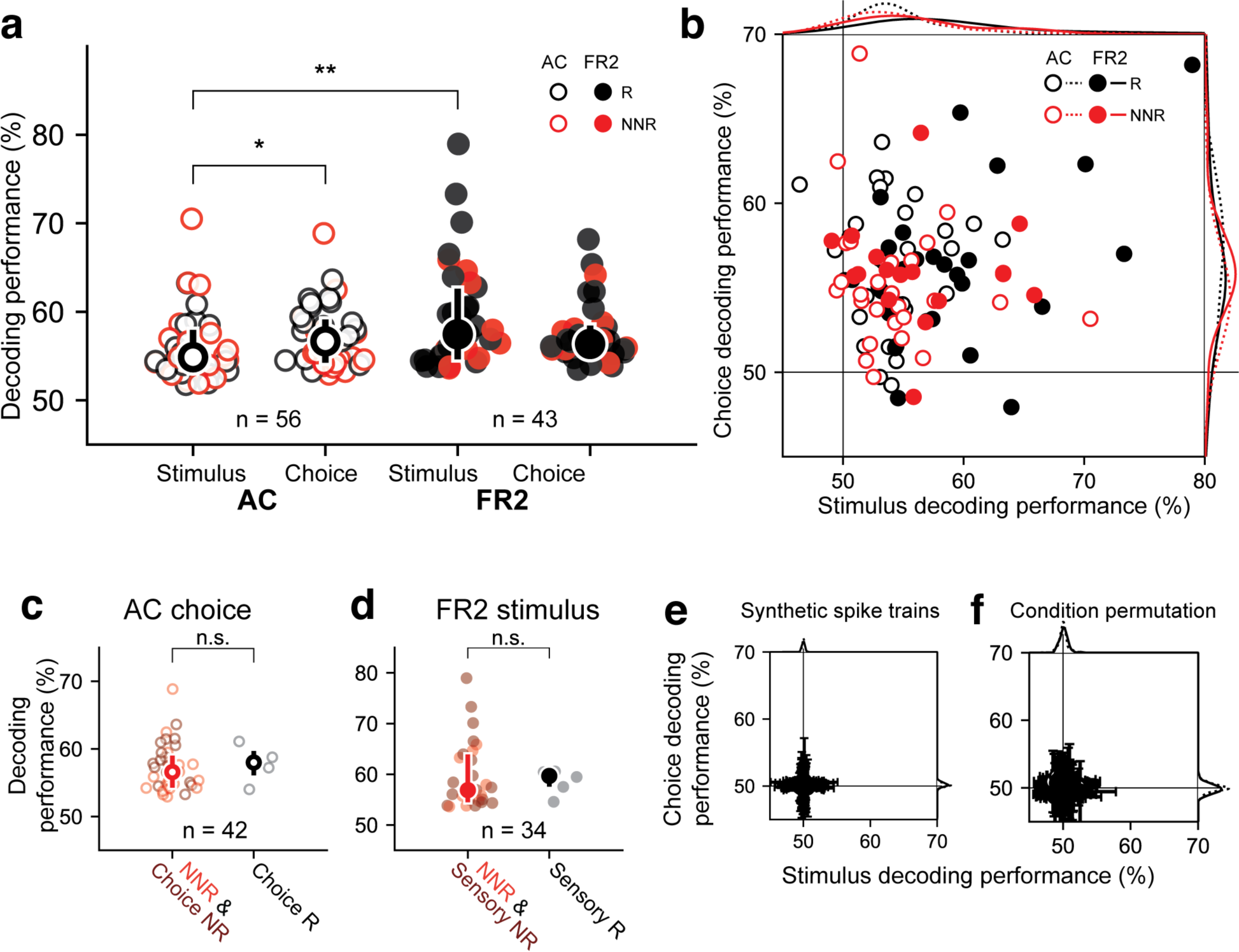
Decoding performance of single-units recorded from AC or FR2. **a**. Decoding performance of single-units for stimulus category and behavioral choice in AC (open circles) and FR2 (filled circles) restricted to those statistically significant relative to synthetically-generated spike trains (p<0.05, permutation test, two-sided). Central symbol with error bars represents group medians and top and bottom quartiles (*p=0.02, **p=0.001, Mann-Whitney U test, two-sided). Black symbols, responsive cells; red symbols, nominally non-responsive cells. **b**. Decoding performance for choice versus stimulus, restricted to those statistically significant relative to synthetically-generated spike trains for either stimulus, choice, or both (p<0.05, permutation test, two-sided). Black symbols, responsive cells; red symbols, nominally non-responsive cells. **c**. Choice decoding performance in AC of nominally non-responsive cells (red) and choice non-responsive (dark-red) versus choice responsive cells (black; i.e. ramping cells). Decoding performance was not statistically different (p=0.32 Mann-Whitney U test, two-sided). Central symbol with error bars represents group medians and top and bottom quartiles. **d**. Stimulus decoding performance in FR2 for nominally non-responsive cells (red) and sensory non-responsive (dark-red) versus choice responsive cells (black; i.e. ramping cells). Decoding performance was not statistically different (p=0.29, Mann-Whitney U test, two-sided). Central symbol with error bars represents group medians and top and bottom quartiles. **e**. Decoding performance for choice versus stimulus, applied to spike trains synthetically generated from sampling (with replacement) over all ISIs observed without regard to stimulus category or behavioral choice. Black, responsive cells; red, nominally non-responsive cells. Error bars represent standard deviation. **f**. Decoding performance for choice versus stimulus, applied to spike trains left intact but trial conditions (stimulus category and behavioral choice) were randomly permuted (1000 permutations per unit). Error bars represent standard deviation.

### Nominally non-responsive cells reveal hidden task information

Can we uncover hidden task information from nominally non-responsive cells? We found that nominally non-responsive cells in both AC and FR2 provided significant information about each task variable (**Fig. 4a,b**, red). The ability to decode was independent of average firing rate (**Supplementary Fig. 9a-f**, 0.30 < r < 0.46), z-score (**Supplementary Fig. 9g-i**, −0.05 < r < 0.05), and ramping activity (**Supplementary Fig. 9j**, −0.02 < r < 0.28). Stimulus decoding performance was also independent of receptive field properties including best frequency and tuning curve bandwidth for AC neurons (**Supplementary Fig. 10**).

We also observed that task information was distributed across both AC and FR2, and neural spike trains from individual units were multiplexed in that they often encoded information about both stimulus category and choice simultaneously (**Fig. 4b**). To establish that multiplexing was not simply a byproduct of the correlation between stimulus and choice variables, an independent measure of multiplexing relying on multiple regression was applied (**Supplementary Fig. 11**). Despite the broad sharing of information about behavioral conditions, there were notable systematic differences between AC and FR2. Surprisingly, neurons in FR2 were more informative about stimulus category than AC, and AC neurons were more informative about choice than stimulus category (**Fig. 4a**, p_ac_=0.016, p_stim_=0.0013, Mann-Whitney U test, two-sided). Both of these observations would not have been detected at the level of the PSTH, as most cells in AC were non-responsive for behavioral choice (no ramping activity, 91/103), yet our decoder revealed that these same cells were as informative as choice responsive cells (**Fig. 4c**, p=0.32 Mann-Whitney U test, two-sided; red circles indicate cells non-responsive for both variables, dark-red cells are choice non-responsive, and black cells are responsive). Similarly, most cells in FR2 were sensory non-responsive (not tone modulated, 60/74), yet contained comparable stimulus information to sensory responsive cells (**Fig. 4d**, p=0.29 Mann-Whitney U test, two-sided; red cells are non-responsive for both variables, dark-red cells are sensory non-responsive, black cells are responsive).

To assess the statistical significance of these results, we tested our algorithm on two shuffled data sets. First, we ran our analysis using synthetically-generated trials that preserved trial length but randomly sampled ISIs with replacement from those observed during a session without regard to condition (**Fig. 4e**). Second, we left trial activity intact, but permuted the stimulus category and choice for each trial (**Fig. 4f**). We restricted analysis to cells with decoding performance significantly different from synthetic spike trains (all cells in **Fig. 4a-d** significantly different from synthetic condition shown in **Fig. 4e**, p<0.05, bootstrapped 1240 times).

To directly assess the extent to which information captured by the ISI distributions in our data set was distinct from the time-varying rate, we compared the performance from our ISI-based decoder to a conventional rate-modulated (inhomogeneous) Poisson decoder^31^ which assumes that spikes are produced randomly with an instantaneous probability equal to the time-varying firing rate. As our model cells illustrate (**Fig. 2**), it is possible to decode using the ISI distributions even when firing rates are uninformative (**Supplementary Fig. 12a**). When applied to our dataset, the ISI-based decoder generally outperformed this conventional rate-based decoder confirming that ISIs capture information distinct from that of the firing rate (**Supplementary Fig. 12b**; Overall stimulus decoding performance: p_ac_=0.0001, p_fr2_=8×10^−6^ Overall choice decoding performance p_ac_=0.0001, p_fr2_=0.02, Mann-Whitney U test, two-sided). Moreover, comparing single trial decoding outcomes demonstrated weak to no correlations between the ISI-based decoder and the conventional rate decoder, further underscoring that these two methods rely on different features of the spike train to decode (**Supplementary Fig. 12c**; stimulus medians: AC=0.10 FR2=0.11; choice medians: AC=0.07, FR2=0.08).

We hypothesize that ISI-based decoding is biologically plausible. Short-term synaptic plasticity and synaptic integration provide powerful mechanisms for differential and specific spike-timing-based coding. We illustrated this capacity by making whole-cell recordings from AC neurons in vivo and in brain slices (**Supplementary Fig. 13a,b**), as well as in FR2 brain slices (**Supplementary Fig. 13c**). In each case, different cells could have distinct response profiles to the same input pattern, with similar overall rates but different spike timings.

Moreover, we note that this type of coding scheme requires few assumptions about implementation, and does not require additional separate integrative processes to compute rates or form generative models. Thus ISI-based decoding coding could be generally applicable across brain areas, as demonstrated here for AC and FR2.

### Nominally non-responsive cells reveal hidden selection rule information in a novel task-switching paradigm

To further demonstrate the generalizability and utility of our approach, we applied our decoding algorithm to neurons that were found to be non-responsive in a previously published study^5^. In this study, rats were trained on a novel auditory stimulus selection task where depending on the context animals had to respond to one of two cues while ignoring the other. Rats were presented with two simultaneous sounds (a white noise burst and a warble). In the “localization” context the animal was trained to ignore the warble and respond to the location of the white noise burst and in the “pitch” context it was trained to ignore the location of the white noise burst and respond to the pitch of the warble (**Fig. 5a**). The main finding of the study is that the pre-stimulus activity in both primary auditory cortex and prefrontal cortex encodes the selection rule (i.e. activity reflects whether the animal is in the localization or pitch context). This conclusion was entirely based on a difference in pre-stimulus firing rate between the two contexts. The authors reported, but did not further analyze, cells that did not modulate their pre-stimulus firing rate. In our nomenclature these cells are “non-responsive for the selection rule”. Using our algorithm we found that the ISI distributions of these cells encoded the selection rule and were significantly more informative than the responsive cells (**Fig. 5b**, p_ac_=5×10^−6^, p_fr2_<0.0002, Mann-Whitney U test, two-sided). This surprising result demonstrates that our algorithm generalizes to novel datasets, and may be used to uncover coding for cognitive variables that are hidden from conventional trial-averaged analyses. Furthermore, these results indicate that as task complexity increases nominally non-responsive cells are differentially recruited for successful task execution.

**Figure 5.**
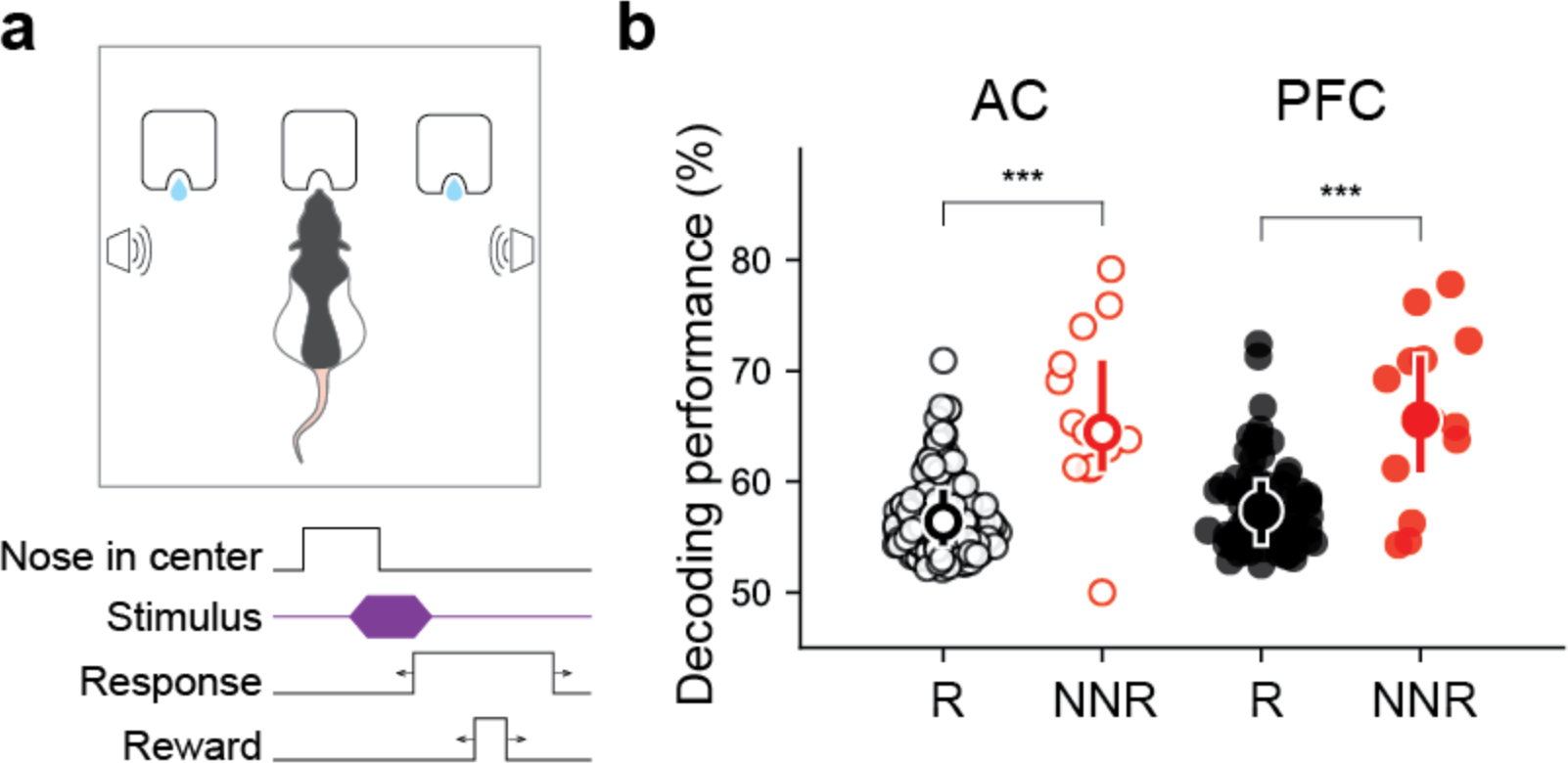
Nominally non-responsive cells in both auditory cortex and prefrontal cortex (PFC) encode selection rule better than responsive cells. **a**. Schematic of novel auditory stimulus selection task. Animals were presented with two simultaneous tones (a white noise burst and warble) and trained to respond to the location of the sound in the “localization” context while ignoring pitch and respond to the pitch while ignoring the location in the “pitch” context (figure adapted from Rodgers & DeWeese 2014, *Neuron).* **b**. Decoding performance for nominally non-responsive cells (similar pre-stimulus firing rates for both pitch and localization blocks) in primary auditory cortex and prefrontal cortex previously reported but not further analyzed in this study (red cells), responsive cells in black (***p_ac_=5×10^−6^, ***p_fr2_<0.0002, Mann-Whitney U test, two-sided).

### Nominally non-responsive ensembles are better predictors of behavioral errors

Downstream brain regions must integrate the activity of many neurons and this ISI-based approach naturally extends to simultaneously recorded ensembles. We therefore asked whether using small ensembles would change or improve decoding. To decode from ensembles, likelihood functions from each cell were calculated independently as before, but were used to simultaneously update the task condition probabilities (p(target | ISI) and p(go | ISI)) on each trial (**Fig. 6a**). Analyzing ensembles of 2-8 neurons in AC and FR2 significantly improved decoding for both variables in FR2 and stimulus decoding in AC (**Fig. 6b**, p_ac stim_=0.04, p_fr2_ stim=1×10^−5^, p_ac_=0.29, p_fr2_ choice=7×10^−5^, Mann-Whitney U test, two-sided). This was not a trivial consequence of using more neurons, as the information provided by individual ISIs on single trials can be contradictory (e.g., compare LLR functions in **Fig. 3c** and **Supplementary Fig. 7c** for 50 ms < ISIs < 120 ms). For ensemble decoding to improve upon single neuron decoding, the ISIs of each member of the ensemble must indicate the same task variable.

**Figure 6.**
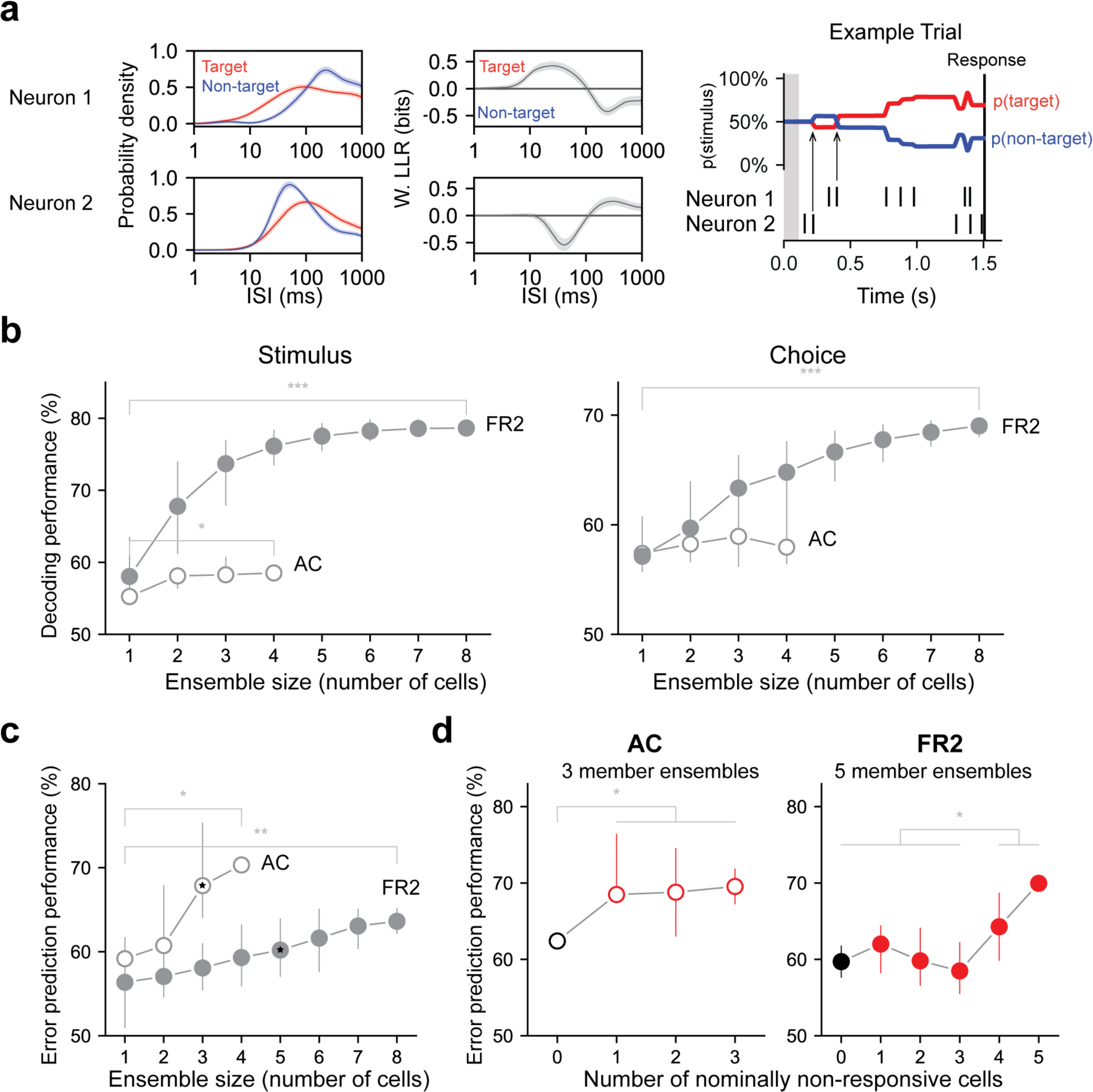
Decoding performance of neuronal ensembles recorded in AC or FR2. **a**. Schematic of ensemble decoding. Left, conditional ISI distributions and corresponding weighted LLR shown for two simultaneously recorded neurons. Right, an example trial where each neuron’s ISIs and LLRs are used to independently update stimulus category according to Bayes’ rule. Arrows indicate the first updates from each neuron. **b**. Stimulus and choice decoding performance for ensembles in AC and FR2 for ensembles of increasing size (Comparing smallest with largest ensembles Stimulus: *p_ac_, ***p_fr2_=1×10^−5^, choice: p_ac_=0.29, ***p_fr2_=7×10^−5^, Mann-Whitney U test, two-sided) c. Error prediction performance in AC and FR2 as a function of ensemble size (*p_ac_=0.03, ***p_fr2_=0.002; comparison between AC and FR2, for 3-member ensembles: p=1.2×10^−5^, for 4-member ensembles: p=0.03, Mann-Whitney U test, two-sided) d. Error prediction performance in AC and FR2 as a function of the number of non-responsive cells in the ensemble (*p_ac_=0.013, ***p_fr2_=0.046 Welch’s t-test), 3 and 5 member ensembles in c. shown for AC and FR2 respectively.

Can our decoding method predict errors on a trial-by-trial basis? In general, trial-averaged PSTHs did not reveal systematic differences between correct and error trials. However, when we examined single-trial performance with our algorithm, ensembles of neurons in AC and FR2 predicted behavioral errors (**Fig. 6c**). In general, ensembles in AC predicted behavioral errors significantly better than those in FR2 (**Fig. 6c**, for 3-member ensembles: p=1.2×10^−5^, for 4-member ensembles: p=0.03, Mann-Whitney U test, two-sided). Interestingly, decoding with an increasing number of nominally non-responsive cells improved error prediction in both AC and FR2 (**Fig. 6d**,( p_ac_ =0.013, p_fr2_=0.046, Welch’s t-test).

### Recurrent neural network model demonstrates nominally non-responsive cells are necessary for task performance and synergistically interact with responsive cells

Our data indicate that nominally non-responsive activity encodes hidden task-information and can predict behavioral errors, but how does it impact task performance? Examining this question requires network modeling to further explore the dynamics of responsive and non-responsive cells. We carried out a series of simulated perturbation experiments on a recurrent neural network trained to perform our frequency recognition task (**Fig. 7a,b**). Inactivation of both responsive and non-responsive units impaired task performance (**Fig. 7c**) indicating that both sub-populations are necessary to complete the task. Surprisingly, inactivation of a random subset of units (including both responsive and non-responsive units) resulted in larger performance decreases than what would be predicted from the psychometric inactivation curves of responsive and non-responsive units treated independently (**Fig. 7d**). This finding suggests that responsive and non-responsive units have a synergistic effect on overall task performance and that both sub-populations should be considered in concert to fully understand behavioral performance.

**Figure 7.**
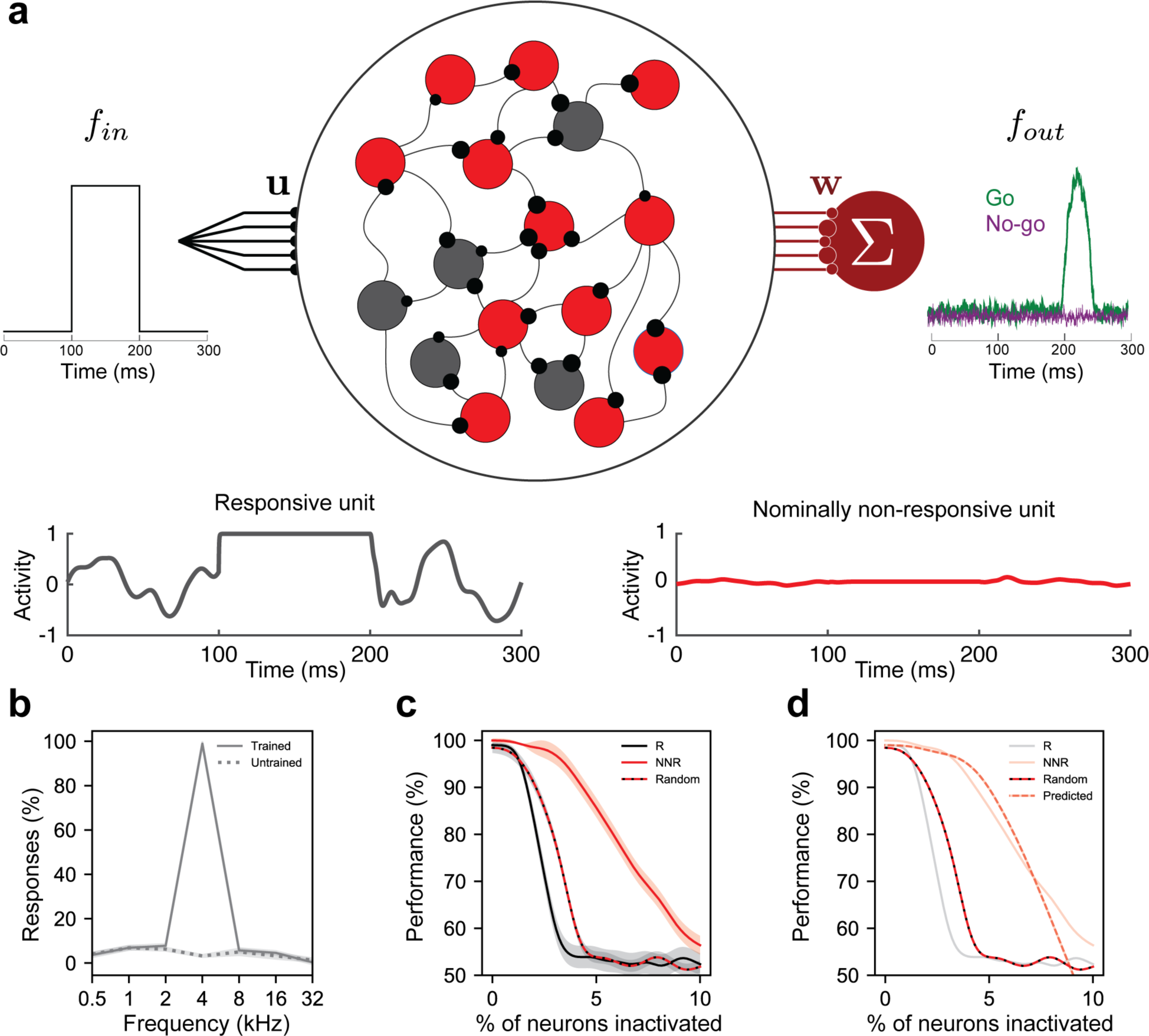
Recurrent neural network model demonstrates nominally non-responsive cells are necessary for task performance and synergistically interact with responsive cells. **a**. Schematic of network design and task details. A recurrent network of model neurons (N = 1000 units, shown here with fewer units for illustration purposes) was trained to produce a choice only in response to a target input (go condition) and no response to non-target inputs (no-go condition). 10% of model neurons (100 units) received the “auditory” input on each 300ms trial, either a target or a nontarget tone presented from 100 - 200 ms (left panel, *f*_*in*_) with weights (**u**), which were generated randomly and left fixed during training. Training was done by activity-dependent modification of the recurrent synaptic weights (black) between responsive units only (R, in gray) and the readout weights (**w**, maroon) from all network units. The overall behavioral choice is read out from the entire network using a readout unit (maroon, *fout). Bottom panel*, activity of an example responsive and non-responsive unit on a single trial. **b**. Task performance of trained versus untrained network. Once trained for 500 steps, these networks were tested for a further 1000 steps or simulated “trials.” Trained networks demonstrated reliable responses only to the target tone (solid line and shading indicate mean ± standard deviation for 10 instantiations of the network). For comparison, untrained networks (dotted line and shading indicate mean ± standard deviation) operating under the same stimulation conditions (run for 1000 trials without training) did not respond to any of the presented “tone” inputs. **c**. Psychometric inactivation curves representing task performance as a function of the number of inactivated responsive cells (black), nominally non-responsive cells (red), and a random subset (dashed red and black line). After training, different fractions of units (between 0 and 10% of units in N=1000 networks, in 0.5% increments) were selected randomly for inactivation from each subpopulation or the entire network *(i.e.*, their currents were set to 0). These partially inactivated networks were then tested for an additional 1000 trials. Data collected from 10 network instantiations (solid line and shading indicate mean ± standard deviation). **d**. Comparison of the psychometric inactivation curve for random subsets (dashed red and black line) with the predicted curve derived from the responsive and non-responsive curves treated independently (dashed red and white line).

### Ensemble consensus-building dynamics underlie hidden task information

While improvements were seen in decoding performance with increasing ensemble size, the ISI distributions/ISI-based likelihood functions were highly variable across individual ensemble members. Thus, we wondered if there was hidden task-related structure in the population activity that evolved over the course of the trial to instantiate behavior. To answer this question, we examined whether local ensembles share the same representation of task variables over the course of the trial. Do they “reach consensus” on how to represent task variables using the ISI (**Fig. 8a**)? Without consensus, a downstream area would need to interpret ensemble activity using multiple disparate representations rather than one unified code (**Fig. 8b**). The firing rates and ISI distributions of simultaneously-recorded units were generally variable across cells requiring an exploratory approach to answer this question (**Fig. 8c**, example three-member ensemble with heterogeneous conditional ISI distributions). Therefore, we examined changes in the distributions of ISIs across task conditions, asking how the moment-to-moment changes in the log-likelihood ratio (LLR) of each cell were coordinated to encode task variables (**Fig. 8c**). We focused on the LLR because it quantifies how the ISI represents task variables for a given cell and summarizes all spike timing information needed by our algorithm (or a hypothetical downstream cell) to decode.

**Figure 8.**
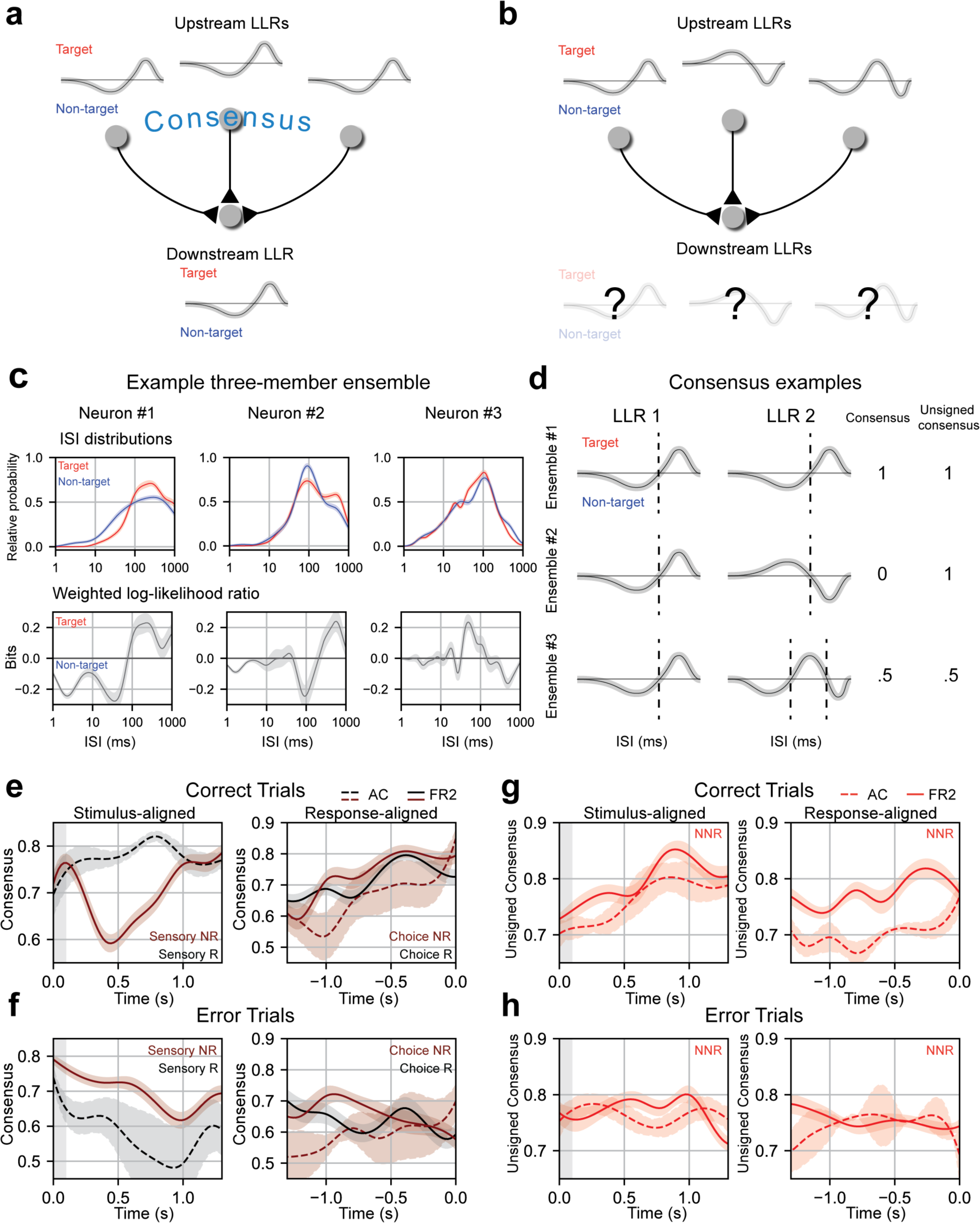
Ensemble consensus-building during behavior. a. Schematic of consensus building in a three-member ensemble. When the LLRs of ensemble members are similar the meaning of any ISI is unambiguous to a downstream neuron. b. Schematic of a three-member ensemble without consensus. The meaning of an ISI depends on the upstream neuron it originates from c. ISI distributions, and LLRs for three members of a sample ensemble. Note that despite differences in ISI distributions, neuron #1 and neuron #2 have similar weighted log-likelihood ratios (ISIs > 200 ms indicate target, ISIs < 200 ms indicate non-target). d. Consensus values for three illustrative two-member ensembles. Ensemble 1 members have identical LLRs, agreeing on the meaning of all ISIs (consensus = 1) and on how the ISIs should be partitioned (unsigned consensus = 1). Ensemble 2 contains cells with LLRs where the ISI meanings are reversed, disagreeing on meaning of the ISIs (consensus = 0) but still agree on how the ISIs should be partitioned (unsigned consensus = 1). Ensemble 3 contains two cells with moderate agreement about the ISI meanings and partitioning, leading to intermediate consensus and unsigned consensus values (0.5 for each). e. Left, mean consensus as a function of time from tone onset (stimulus-aligned) on correct trials for three-member sensory responsive ensembles in AC (two or more members sensory responsive; black dotted line; n=11 ensembles) and sensory non-responsive ensembles in FR2 (two or more members sensory non-responsive; dark red solid line; n=101 ensembles). Standard deviation shown around each mean trendline. FR2 sensory non-responsive cells consistently reached consensus and then diverged immediately after stimulus presentation (Δconsensus, *t* = 0 to 0.42 s, p_snr_ = 1.3 ×10^−4^ Wilcoxon test, two-sided). AC responsive ensembles (black) increase consensus until 750 ms Δconsensus, *t* = 0 to 0.81 s, psR = 0.046 Wilcoxon test, two-sided). Right, mean consensus as a function of time to behavioral response (response-aligned) on correct trials for three-member choice responsive ensembles (two or more members choice responsive; black) in FR2 (solid line; n=47 ensembles) and choice non-responsive (two or more members choice non-responsive; dark red) in AC (dotted line; n=11 ensembles) and FR2 (solid line; n=57 ensembles). Standard deviation shown around each mean trendline. On correct trials, choice responsive (black) and choice non-responsive ensembles (dark red) in both regions reached high consensus values ~500 ms before response (Δconsensus, *t* = −1.0 to 0.0 s, p_cnr_ = 6.6×10^−6^, p_cr_ = 0.041 Wilcoxon test, two-sided). f. As in e, but for error trials (Δconsensus, correct vs. error trials, stimulus: p_snr_= 0.007, p_sr_ = 0.065, choice: pcnr = 0.0048, pcR = 0.065 Mann-Whitney U test, two-sided). g. Unsigned consensus index for nominally non-responsive ensembles (two or more members nominally non-responsive) in AC (dotted line; n=13 ensembles) and FR2 (solid line; n=36 ensembles), stimulus-aligned (left, Δconsensus, *t* = 0 to 0.89 s, p = 1.7×10^−5^ Wilcoxon test, two-sided) and response-aligned (right, Δconsensus, *t* = −1.0 to 0.0 s, p = 0.0011 Wilcoxon test, two-sided). On correct trials, ensembles reach high values of unsigned consensus ~750 ms after tone onset and within 500 ms of behavioral response. h. As in g, but for error trials (Δconsensus, correct vs. error trials, p = 1.9×10^−9^ Mann-Whitney U test, two-sided).

We examined how ensembles coordinate their activity moment-to-moment over the course of the trial by quantifying the similarity of the LLRs across cells in a sliding window. Similarity was assessed by summing the LLRs of ensemble members, calculating the total area underneath the resulting curve, and normalizing this value by the sum of the areas of each individual LLR. We refer to this quantified similarity as ‘ consensus’; a high consensus value indicates that the LLRs generally agree on stimulus or choice while a low value indicates disagreement at that moment (**Fig. 8d**). We should emphasize that successful ensemble decoding (**Fig. 6**) does not require the LLRs of ensemble members to be related in any way; therefore, structured LLR dynamics (**Fig. 8**) are not simply a consequence of how our algorithm is constructed.

While the conventional trial-averaged PSTH of non-responsive ensembles recorded in AC and FR2 showed no task-related modulation, our analysis revealed structured temporal dynamics of the LLRs (captured by the consensus value). In FR2, sensory non-responsive ensembles (ensembles in which at least two out of three cells were not tone-modulated) encode stimulus information using temporally-precise stimulus-related dynamics on correct trials. The stimulus representation of sensory non-responsive ensembles reached consensus rapidly after stimulus onset followed by divergence (**Fig. 8e**, stimulus-aligned, solid line, Δconsensus, *t* = 0 to 0.42 s, = p_snr_=1.3 ×10^−4^ Wilcoxon test, two-sided). Sensory responsive ensembles in AC increased consensus beyond stimulus presentation, reaching a maximum ~750 ms after tone onset on correct trials (**Fig. 8e** stimulus-aligned, dotted line, Δconsensus, *t* = 0 to 0.81 s, p_sr_ = 0.046 Wilcoxon test, two-sided). For choice-related activity, choice non-responsive ensembles in both regions as well as choice responsive ensembles in FR2 each reached consensus within 500 ms of the behavioral response (**Fig. 8e**, response-aligned, Δconsensus, t = −1.0 to 0.0 s, p_cnr_= 6.6× 10^−6^, p_cr_ = 0.041 Wilcoxon test, two-sided). Importantly, this temporally precise pattern of consensus building is not present on error trials (**Fig. 8f**, Δconsensus, correct trials vs. error trials, stimulus:p_snr_ =0.007, p_sr_ = 0.065, choice: p_cnr_= 0.0048, pcR = 0.065 Mann-Whitney U test, two-sided).

These results reveal that consensus-building and divergence occur at key moments during the trial for successful execution of behavior in a manner that is invisible at the level of the PSTH. As sensory and choice non-responsive ensembles participated in these dynamics, changes in the consensus value cannot simply be a byproduct of correlated firing rate modulation due to tone-evoked responses or ramping. While consensus-building can only indicate a shared representation, divergence can indicate one of two things: (1) the LLRs of each cell within an ensemble are completely dissimilar or (2) they are ‘out of phase’ with one another - the LLRs partition the ISIs the same way (**Fig. 8d**, dotted lines), but the same ISIs code for opposite behavioral variables. This distinction is important because (2) implies coordinated structure of ensemble activity (the partitions of the ISI align) whereas (1) does not. To distinguish between these two possibilities we used the ‘unsigned consensus’, a second measure sensitive to the ISI partitions but insensitive to the sign of the LLR. Both ‘in phase’ and perfectly ‘out of phase’ LLRs would produce an unsigned consensus of 1 whereas unrelated LLRs would be closer to 0 (**Fig. 8d**). For example, in the second row of **Figure 8d**, both cells agree that ISIs < 100 ms indicate one stimulus category and ISIs > 100 ms indicate another, but they disagree about which set of ISIs mean target and which mean non-target. This results in a consensus value of 0 (out of phase) but an unsigned consensus value of 1.

Using this metric, we found that the unsigned consensus pattern for nominally non-responsive ensembles (ensembles with two or more nominally non-responsive members) were shared between AC and FR2 - increasing until ~750 ms after tone onset on correct trials (**Fig. 8g**, stimulus-aligned, Δconsensus, *t* = 0 to 0.89 s, p = 1.7× 10^−5^ Wilcoxon test, two-sided). This intriguing observation reveals the timed coordination of nominally non-responsive ensembles coincident across brain regions and suggests that these cells may constitute a distinct functional network separate from that of responsive cells. Nominally non-responsive ensembles in AC and FR2 also increased their unsigned consensus immediately before behavioral response (although values in AC were lower overall; **Fig. 8g**, response-aligned, Δconsensus, *t* = −1.0 to 0.0 s, p = 0.0011 Wilcoxon test, two-sided). This pattern of consensus-building was only present on correct trials but not incorrect trials (**Fig. 8h**, Δconsensus compared to error trials, p = 1.9× 10^−9^ Mann-Whitney U test, two-sided) suggesting that behavioral errors might result from a general lack of consensus between ensemble members. In summary, we have shown that cells which appear unmodulated during behavior do not encode task information independently, but do so by synchronizing their representation of behavioral variables dynamically during the trial.

## Discussion

Using a straightforward, single-trial, ISI decoding algorithm that makes few assumptions about the proper model for neural activity, we found task-specific information extensively represented by what appeared to be nominally non-responsive neurons in both AC and FR2. Furthermore, the degree to which single neurons were task-modulated was uncorrelated with conventional response properties including frequency tuning. AC and FR2 each represent both task-variables; furthermore, in both regions we identified many multiplexed neurons that simultaneously represented the sensory input and the upcoming behavioral choice including non-responsive cells. This highlights that the cortical circuits that generate behavior exist in a distributed network – blurring the traditional modular view of sensory and frontal cortical regions.

Most notably, FR2 has a better representation of task-relevant auditory stimuli than AC. The prevalence of stimulus information in FR2 might be surprising given that AC reliably responds to pure tones in untrained animals; however, when tones take on behavioral significance, this information is encoded more robustly in frontal cortex, suggesting that this region is critical for identifying the appropriate sensory-motor association. Furthermore, the stark improvement in stimulus encoding for small ensembles in FR2 suggests that task-relevant stimulus information is reflected more homogeneously in local firing activity across FR2 (perhaps through large scale ensemble consensus-building) while this information is reflected in a more complex and distributed manner throughout AC.

The finding that the ISI-based approach of our algorithm is not reducible to rate despite their close mathematical relationship raises the question of how downstream regions could respond preferentially to specific ISIs. Our whole cell recordings from both AC and FR2 demonstrate that different postsynaptic cells can respond differently to the same input pattern with a fixed overall rate emphasizing the importance of considering a code sensitive to precise spike-timing (**Supplementary Fig. 13**). Furthermore, this is supported by experimental and theoretical work showing that single neurons can act as resonators tuned to a certain periodicity of firing input^32^. This view could also be expanded to larger neuronal populations comprised of feedback loops that would resonate in response to particular ISIs. In this case, cholinergic neuromodulation could offer a mechanism for adjusting the sensitivities of such a network during behavior on short time-scales by providing rapid phasic signals^33^.

It is still unclear what the relevant timescales of decoding might be in relation to phenomena such as membrane time constants, periods of oscillatory activity, and behavioral timescales. Given that our ISI-based decoder and conventional rate-modulated decoders reveal distinct information, future approaches might hybridize these rate-based and temporal-based decoding methods to span multiple timescales.

We have also shown that underlying the task-relevant information encoded by each ensemble is a rich set of consensus-building dynamics that is invisible at the level of the PSTH. Ensembles in both FR2 and AC underwent stimulus and choice-related consensus building that was only observed when the animal correctly executed the task. Moreover, nominally non-responsive cells demonstrated temporal dynamics synchronized across regions which were distinct from responsive ensembles. This raises the possibility that nominally non-responsive ensembles constitute a discrete functional network distinguishable from responsive ensembles. These results underscore the importance of measuring neural activity in behaving animals and using unbiased and generally-applicable analytical methods, as the response properties of cortical neurons in a behavioral context become complex in ways that challenge our conventional assumptions^7–9,34^.

## Methods

### Behavior

All animal procedures were performed in accordance with National Institutes of Health standards and were conducted under a protocol approved by the New York University School of Medicine Institutional Animal Care and Use Committee. We used 23 adult Sprague-Dawley male and female rats (Charles River) in the behavioral studies. Animals were food restricted and kept at 85% of their initial body weight, and maintained at a 12 hr light/12 hr dark cycle.

Animals were trained on a go/no-go audiomotor task^8,13^. Operant conditioning was performed within 12” L × 10” W × 10.5” H test chambers with stainless steel floors and clear polycarbonate walls (Med Associates), enclosed in a sound attenuation cubicle and lined with soundproofing acoustic foam (Med Associates). The nose and reward ports were both arranged on one of the walls with the speaker on the opposite wall. The nose port, reward port, and the speaker were controlled and monitored with a custom-programmed microcontroller. Nose port entries were detected with an infrared beam break detector. Auditory stimuli were delivered through an electromagnetic dynamic speaker (Med Associates) calibrated using a pressure field microphone (ACO Pacific).

Animals were rewarded with food for nose poking within 2.5 seconds of presentation of the target tone (4 kHz) and given a short 7-second time-out for incorrectly responding to non-target tones (0.5, 1, 2, 8, 16, 32 kHz). Incorrect responses include either failure to enter the nose port after target tone presentation (miss trials) or entering the nose port after non-target tone presentation (false alarms). Tones were 100 msec in duration and sound intensity was set to 70 dB SPL. Tones were presented randomly with equal probability such that each stimulus category was presented. The inter-trial interval delays used were 5, 6, 7, or 8 seconds.

For experiments involving muscimol, we implanted bilateral cannulas in either FR2 (+2.0 to +4.0 mm AP, ±1.3 mm ML from Bregma) of 7 animals or AC (−5.0 to −5.8 mm AP, 6.5-7.0 mm ML from Bregma) of 3 animals. We infused 1 μL of muscimol per side into FR2 or infused 2 μL of muscimol per side into AC, at a concentration of 1 mg/mL. For saline controls, equivalent volumes of saline were infused in each region. Behavioral testing was performed 30-60 minutes after infusions. Power analysis was performed to determine sample size for statistical significance with a power of β: 0.8; these studies required at least 3 animals, satisfied in the experiments of **Supplementary Fig. 3b,e**. For motor control study, animals could freely nose poke for food reward without presentation of auditory stimuli after muscimol and saline infusion.

### Implant preparation and surgery

Animals were implanted with microdrive arrays (Versadrive-8 Neuralynx) in either AC (8 animals) or FR2 (7 animals) after reaching behavioral criteria of d’ ≥ 1.0. For surgery, animals were anesthetized with ketamine (40 mg/kg) and dexmedetomidine (0.125 mg/kg). Stainless steel screws and dental cement were used to secure the microdrive to the skull, and one screw was used as ground. Each drive consisted of 8 independently adjustable tetrodes. The tetrodes were made by twisting and fusing four polyimide-coated nichrome wires (Sandvik Kanthal HP Reid Precision Fine Tetrode Wire; wire diameter 12.5 μm). The tip of each tetrode was gold-plated to an impedance of 300-400 kOhms at 1 kHz (NanoZ, Neuralynx).

### Electrophysiological recordings & unit isolation

Recordings in behaving rats were performed as previously described^8^. After the animal recovered from surgery (~7 days) recordings began once performance returned to presurgery levels. Tetrodes were advanced ~60 μm 12 hours prior to each recording session, to a maximum of 2.5mm (for FR2) or 2.0 mm (for AC) from the pial surface. For recording, signals were first amplified onboard using a small 16-bit unity-gain preamplifier array (CerePlex M, Blackrock Microsystems) before reaching the acquisition system. Spikes were sampled at 30 kS/sec and bandpass filtered between 250 Hz and 5 kHz. Data were digitized and all above-threshold events with signal to noise ratios > 3:1 were stored for offline spike sorting. Single-units were identified on each tetrode using OfflineSorter (Plexon Inc.) by manually classifying spikes projected as points in 2D or 3D feature space. The parameters used for sorting included the waveforms projection onto the first two principal components, energy, and nonlinear energy. Artifacts were rejected based on refractory period violations (< 1 msec). Clustering quality was assessed based on the Isolation Distance and Lratio sorting quality metrics. To be initially included for analysis, cells had to have > 3 spikes per trial for 80% of trials to ensure that there were enough ISIs to reliably estimate the ISI probability density functions.

### Statistical tests for non-responsiveness

We used two positive statistical tests for non-responsiveness: one to establish a lack of tone-modulation, the other to establish a lack of ramping activity. To accommodate the possibility of tone onset and offset responses, we performed our tone-modulation test on a 100 ms long tone presentation window as well as the 100 ms window immediately after tone presentation. The test compared the number of spikes during each of these windows to inter-trial baseline activity as measured by three sequential 100 ms windows preceding tone onset. Three windows were chosen to account for variability in spontaneous spike counts. Given that spike counts are discrete, bounded, and non-normal, we used bootstrapping to evaluate whether the mean change in spikes during tone presentation was sufficiently close to zero (in our case 0.1 spikes). We subsampled 90% of the spike count changes from baseline, calculated the mean of these values, and repeated this process 5000 times to construct an empirical distribution of means. If 95% of the subsampled means values were between −0.1 and 0.1 we considered the cell sensory non-responsive (p<0.05). The value of 0.1 spikes was chosen to be conservative as it is equivalent to an expected change of 1 spike every 10 trials. This is a conservative, rigorous method for establishing sensory non-responsiveness that is commensurate with more standard approaches for establishing tone responsiveness such as the z-score.

To quantify the observed sustained increase in firing rate preceding the behavioral response a ramp index was calculated adapted from the ‘build-up rate’ used in previous literature^31^. First, the trial averaged firing rate was determined in 50 msec bins leading up to the behavioral response. We then calculated the slope of a linear regression in a 500 msec long sliding window beginning 850 msec before behavioral response. The maximum value of these slopes was used as the ‘ramp index’ for each cell. Cells were classified as choice non-responsive if the ramp index did not indicate an appreciable change in the firing rate (less than 50% change) established via bootstrapping. Cells that were shown to be both sensory and choice non-responsive were considered nominally non-responsive overall (**Fig. 4a,b**, red circles).

### Additional firing statistics

Spontaneous average firing rate was established by averaging spikes in a 100 msec time window immediately prior to tone onset on each trial. To quantify tone modulated responses observed during stimulus presentation, we calculated z-scores of changes in spike count from 100 msec before tone onset to 100 msec during tone presentation:

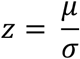

where *μ* is the mean change in spike count and *σ* is the standard deviation of the change in spike count.

### Analysis of receptive field properties

Receptive fields were constructed by calculating the average change in firing rate from 50 ms before tone onset to 50 ms during tone presentation. The window used during tone presentation was identical to that used to calculate the z-score. Best frequency was defined as the frequency where the largest positive deviation in the evoked firing rate was observed. Tuning curve bandwidth was determined by calculating the width of the tuning curve measured at the mean of the maximum and minimum observed evoked firing rates.

### In vivo whole-cell recordings

Sprague-Dawley rats 3-5 months old were anesthetized with pentobarbital. Experiments were carried out in a sound-attenuating chamber. Series of pure tones (70 dB SPL, 0.5-32 kHz, 50 msec, 3 msec cosine on/off ramps, inter-tone intervals between 50-500 msec) were delivered in pseudo-random sequence. Primary AC location was determined by mapping multiunit responses 500-700 μm below the surface using tungsten electrodes. In vivo whole-cell voltage-clamp recordings were then obtained from neurons located 400-1100 μm below the pial surface. Recordings were made with an AxoClamp 2B (Molecular Devices). Whole-cell pipettes (5-9 MΩ) contained (in mM): 125 Cs-gluconate, 5 TEACl, 4 MgATP, 0.3 GTP, 10 phosphocreatine, 10 HEPES, 0.5 EGTA, 3.5 QX-314, 2 CsCl, pH 7.2. Data were filtered at 2 kHz, digitized at 10 kHz, and analyzed with Clampfit 10 (Molecular Devices). Tone-evoked excitatory postsynaptic currents were recorded at −70 mV.

### In vitro whole-cell recordings

Acute brain slices of AC or FR2 were prepared from 2-5 month old Sprague-Dawley rats. Animals were deeply anesthetized with a 1:1 ketamine/xylazine cocktail and decapitated. The brain was rapidly placed in ice-cold dissection buffer containing (in mM): 87 NaCl, 75 sucrose, 2.5 KCl, 1.25 NaH2PO4, 0.5 CaCl2, 7 MgCl2, 25 NaHCO3, 1.3 ascorbic acid, and 10 dextrose, bubbled with 95%/5% O2/CO2 (pH 7.4). Slices (300-400 μm thick) were prepared with a vibratome (Leica), placed in warm dissection buffer (32-35°C) for 10 min, then transferred to a holding chamber containing artificial cerebrospinal fluid at room temperature (ACSF, in mM: 124 NaCl, 2.5 KCl, 1.5 MgSO4, 1.25 NaH2PO4, 2.5 CaCl2, and 26 NaHCO3,). Slices were kept at room temperature (22-24°C) for at least 30 minutes before use. For experiments, slices were transferred to the recording chamber and perfused (2-2.5 ml min^−1^) with oxygenated ACSF at 33°C. Somatic whole-cell current-clamp recordings were made from layer 5 pyramidal cells with a Multiclamp 700B amplifier (Molecular Devices) using IR-DIC video microscopy (Olympus). Patch pipettes (3-8 MW) were filled with intracellular solution containing (in mM): 120 K-gluconate, 5 NaCl, 10 HEPES, 5 MgATP, 10 phosphocreatine, and 0.3 GTP. Data were filtered at 2 kHz, digitized at 10 kHz, and analyzed with Clampfit 10 (Molecular Devices). Focal extracellular stimulation was applied with a bipolar glass electrode (AMPI Master-9, stimulation strengths of 0.1-10 V for 0.3 msec). Spike trains recorded from AC and FR2 units during behavior were then divided into 150-1000 msec fragments, and used as extracellular input patterns for these recordings.

### ISI-based single-trial Bayesian decoding

Our decoding method was motivated by the following principles: First, single-trial spike timing is one of the only variables available to downstream neurons. Any observations about trial-averaged activity must ultimately be useful for single-trial decoding, in order to have behavioral significance. Second, there may not be obvious structure in the trial-averaged activity to suggest how non-responsive cells participate in behaviorally-important computations. This consideration distinguishes our method from other approaches that rely explicitly or implicitly on the PSTH for interpretation or decoding^4,10,11,35–37^. Third, we required a unified approach capable of decoding from both responsive and non-responsive cells in sensory and frontal areas with potentially different response profiles. Fourth, our model should contain as few parameters as possible to account for all relevant behavioral variables (stimulus category and behavioral choice). This model-free approach also distinguishes our method from others that rely on parametric models of neural activity.

These requirements motivated our use of ISIs to characterize neuronal activity. For non-responsive cells with PSTHs that displayed no systematic changes over trials or between task conditions, the ISI distributions can be variable. The ISI defines spike timing relative to the previous spike and thus does not require reference to an external task variable such as tone onset or behavioral response. In modeling the distribution of ISIs, we use a non-parametric Kernel Density Estimator that avoids assumptions about whether or not firing occurs according to a Poisson (or another) parameterized distribution. We used maximum likelihood to estimate the bandwidth of the Kernel Density Estimator in a data-driven manner. Finally, the use of the ISI was also motivated by previous work demonstrating that the ISI can encode sensory information^28–30^ and that precise spike timing has been shown to be important for sensory processing in rat auditory cortex^38,39^.

*Training probabilistic model:* Individual trials were defined as the time from stimulus onset to the response time of the animal (or average response time in the case of no-go trials). Trials were divided into four categories corresponding to each of the four possible variable combinations (target/go, target/no-go, non-target/go, non-target/no-go). Approximately 90% of each category was set aside as a training set in order to determine the statistical relationship between the ISI and the two task variables (stimulus category, behavioral choice).

Each ISI observed was sorted into libraries according to the stimulus category and behavioral choice of the trial. The continuous probability distribution of finding a particular ISI given the task condition of interest (target or non-target, go or no-go) was then inferred using nonparametric Kernel Density Estimation with a Gaussian kernel of bandwidth set using a 10-fold cross-validation^40^. Because the domain of the distribution of ISIs is by definition positive (ISI > 0), the logarithm of the ISI was used to transform the domain to all real numbers. In the end, we produced four continuous probability distributions quantifying the probability of observing an ISI on a trial of a given type: p(ISI|target), p(ISI|non-target), p(ISI|go), and p(ISI|no-go). These distributions were estimated in a 1 second sliding window starting at the beginning of the trial to account for dynamic changes in the ISI distributions over the course of the trial.

*Decoding:* The remaining 10% of trials in the test set are then decoded using the ISI likelihood function described in the previous section. Each trial begins with agnostic beliefs about the stimulus category and the upcoming behavioral choice (p(target) = p(non-target) = 50%). Each time an ISI was observed, beliefs were updated according to Bayes’ rule with the four probability distributions obtained in the previous section serving as the likelihood function. To update beliefs in the probability of the target tone when a particular ISI has been observed we used the following relationship:

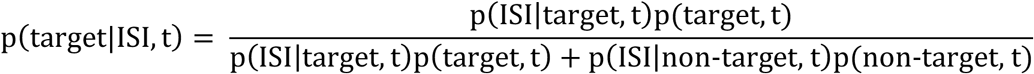

On the left hand side are the updated beliefs about the probability of a target. When the next ISI is observed this value would be inserted as p(target, t) on the right side of the equation and updated once more. Using the probability normalization, p(non-target, t) can be determined,

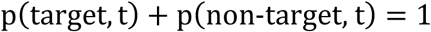

Similarly, for, choice,

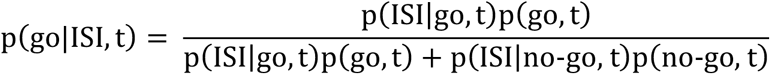

and

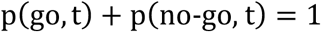

Continuing this process over the course of the trial, we obtain four probabilities - one for each of the variable outcomes - as a function of time during the trial: p(target, t), p(non-target, t), p(go, t), and p(no-go, t). At each moment, the total probability of both stimuli and both choices are 1. The prediction for the entire trial was assessed at the end of the trial. For comparison to choice probability and latency decoding, the certainty was ignored and the preferred variable was taken as the prediction with 100% certainty.

The overall likelihood for a spike train is then simply equal to product of the likelihoods for each ISI observed over the course of the trial,

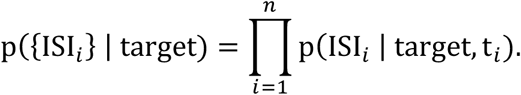

We used 10-fold cross-validation, meaning the trials in the four stimulus categories were randomly divided into ten parts and each part took a turn acting as the test set with the remaining 90% of trials acting as a training set. To estimate the statistical certainty of these results we used bootstrapping with 124 repetitions (except in the case of the null hypotheses where 1240 repetitions were used).

*Ensemble decoding:* Ensemble decoding proceeded very similarly to the single-unit case. The ISI probability distributions for each neuron in the ensemble were calculated independently as described above. However, while decoding a given trial, the spike trains of all neurons in the ensemble were used to simultaneously update the beliefs about stimulus category and behavioral choice. In other words, p(stimulus, t) and p(choice, t) were shared for the entire ensemble but each neuron updated them independently using Bayes’ rule whenever a new ISI was encountered. The joint likelihood of observing a set of ISIs during a trial is then the product of the likelihoods of each neuron independently. For example, for a two neuron ensemble, the combined likelihood, p_12_, of observing the set {ISI_*i*_}_1_= from neuron 1 and {ISI_*i*_}_2_ from neuron 2 is

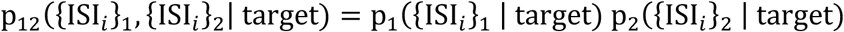

where p_*j*_-is the likelihood of observing a given set of ISIs from neuron *j.*

### Synthetic spike trains

To test the null hypothesis that the ISI-based single-trial Bayesian decoder performance was indistinguishable from chance, synthetic spike trains were constructed for each trial of a given unit by randomly sampling with replacement from the set of all observed ISIs regardless of the original task variable values (synthetic spike trains, **Fig. 4e**). In principle under this condition, ISIs should no longer bear any relationship to the task variables and decoding performance should be close to 50%. For single-unit responses, this randomization was completed 1240 times. Significance from the null was assessed by a direct comparison to the 124 bootstrapped values observed from the true data to the 1240 values observed under the null hypotheses. The p-value was determined as the probability of finding a value from this synthetic condition that produced better decoding performance than the values actually observed as in a standard permutation test.

As a secondary control, we used a traditional permutation test whereby observed spike trains were left intact, but the task variables that correspond to each spike train were randomly permuted (condition permutation, **Fig. 4f**). This process was completed 1240 times.

### Rate-modulated Poisson decoding

To decode using the trial-averaged firing rate, we implemented a standard method^31^ which uses the probability of observing a set of *n* spikes at times *t*_*1*_, …, *t*_*n*_ assuming those spikes were generated by a rate-modulated Poisson process (**Supplementary Fig. 12**). First, we use a training set comprising 90% of trials to estimate the time-varying firing rate for each condition from the PSTH ( *r*_target_(*t*),*r*_non-target_(*t*),*r*_go_(*t*),*r*_no-go_(*t*)) by Kernel Density Estimation with 10-fold cross-validation. The remaining 10% of spike trains are then decoded using the probability of observing each spike train on each condition assuming they were generated according to a rate-modulated Poisson process

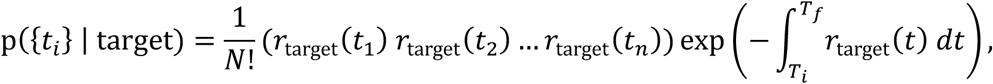

where *T*_*i*_ and *T*_*f*_ are the beginning and end of the trial respectively. This likelihood function is straightforward to interpret: the first product is the probability of observing spikes the spikes at the times they were observed (where the *1/N!* term serves to divide out by the number of permutations of spike labels) and the exponential term represents the probability of silence in the periods between spikes. For comparison with our method, we can reformulate this equation using interspike intervals, if we first break up the exponential integral into domains that span the observed interspike intervals.

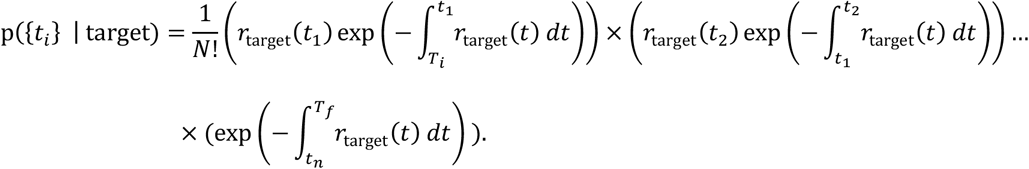

Collecting the first and last terms relating to trial start and trial end as

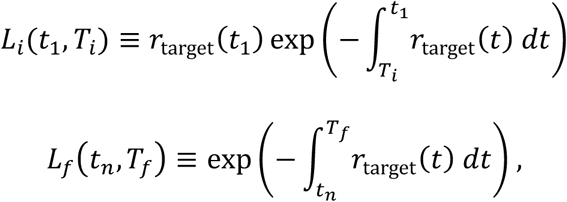

this becomes

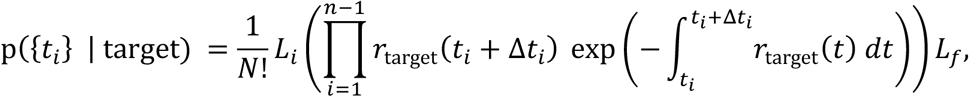

where *Δt*_*i*_ is the time difference between spikes *t*_*i*_ and *t*_*i+1*_ The interpretation of each term in the product is straightforward: it is the infinitesimal probability of observing a spike a time *Δt* after a spike at time *t* multiplied by the probability of observing no spikes in the intervening time. In other words, it is simply p(ISI | target, *t*), the probability of observing an ISI conditioned on observing the first spike at time *t*, as predicted by the assumption of a rate-modulated Poisson process. We can easily verify that this term is normalized which allows us to write,

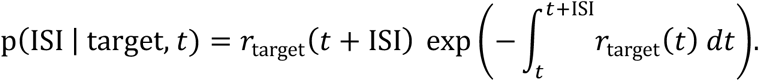

With the exception of the terms relating to trial start and end, we can then view the likelihood of a spike train as resulting from the likelihood of the individual ISIs (just as with our ISI-decoder),

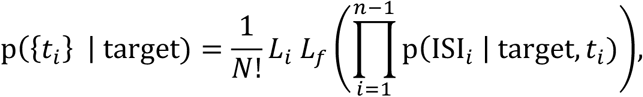

with the key difference that these ISI probabilities are inferred from the firing rate rather than estimated directly using non-parametric methods.

### Inferring the ISI distribution predicted by a rate-modulated Poisson process

To compare the ISI distribution inferred using non-parametric methods to one predicted by a rate-modulated Poisson process we use the relationship above to calculate the predicted probability of observing an ISI of given length within the 1 second window used for our non-parametric estimates. The formula above assumes a spike has already occurred at time t, so we multiply by the probability of observing a spike at time *t*, p(*t* | target) = *r*_target_(*t*), to obtain the total probability of finding an ISI at any given point in the trial.

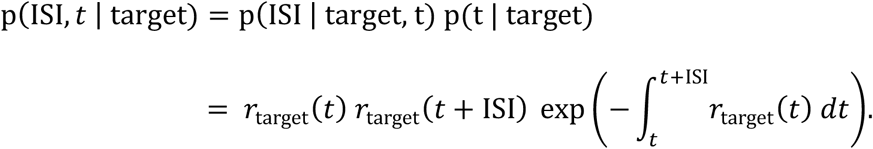

In other words, the probability of observing an ISI beginning at time t is simply the probability of observing spikes at times *t* and *t* + ISI with silence in between.

The probability of observing an ISI at *any* time within a time window spanning *wi* to *wf* is simply the integral of this ISI probability as a function of time across the window. To ensure the final spike occurs before *w*_*f*_ the integral spans *w*_*i*_ to *(w*_*f*_ - ISI),

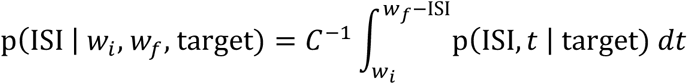

where *C* is a normalization constant which ensures p(ISI | *w_i_, w_f_*, target) integrates to 1,

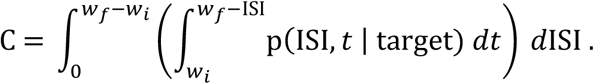

### Regression based method for verifying multiplexing

For each cell, we fit a Logit model for both the stimulus and choice decoding probabilities on individual trials with the true stimulus category and behavioral choice as regressors. We then calculated the extent to which the stimulus decoding probability was determined by true stimulus category by subtracting the regression coefficient for stimulus from that of choice (**Supplementary Fig. 11a**, x-axis, stimulus selectivity index); when this number is positive it indicates that stimulus was a stronger predictor of stimulus decoding on a trial-by-trial basis. The same process was repeated for choice (**Supplementary Fig. 11a**, y-axis, choice selectivity index). According to this analysis we took multiplexed cells to be those that were positive for both measures (**Supplementary Fig. 11a**, orange symbols, 19/90 cells). In other words, multiplexed cells were cells for which stimulus decoding probabilities were primarily a result of true stimulus category *and* choice decoding probabilities were primarily a result of true behavioral choice.

Given the moderate negative correlation for these indices we projected each of these points onto their linear regression to create a one-dimensional regression-based uniplexing index. Cells with a value near zero are the multiplexed cells described above and cells with positive or negative values are primarily stimulus or choice selective (**Supplementary Fig. 11a**).

We compared the uniplexing values produced by this regression method to those produced by examining only the average decoding performance for stimulus and choice (**Supplementary Fig. 11b**). A decoding-based uniplexing index was defined as the difference between average stimulus and choice decoding for each cell. When these two values are comparable this measure returns a value close to zero and the cell is considered multiplexed; moreover, cells that are uniplexed for stimulus or choice receive positive and negative values respectively just as with the regression based measure. While the overall magnitude of these two measures need not be related, both measures of multi/uniplexing rank cells on a one-dimensional axis from choice uniplexed to multiplexed to stimulus uniplexed centered on zero.

### Weighted log likelihood ratio

The log likelihood ratio (LLR) was calculated by first calculating the conditional ISI probabilities and then taking the difference of the logarithm of these distributions. For stimulus,

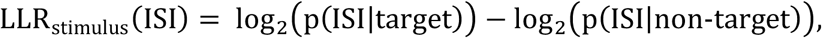

and for choice,

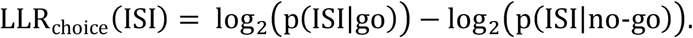

The weighted LLR weights the LLR according to the prevalence of a given ISI. For stimulus,

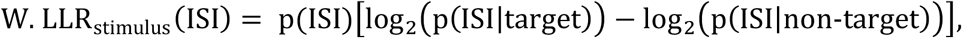

and for choice,

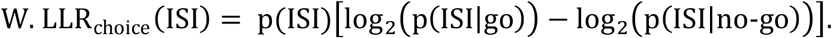

### Consensus and unsigned consensus

The consensus value evaluates the extent to which the LLR (or weighted LLR) is shared across an ensemble. It is the norm of the sum of the LLRs (W. LLRs) divided by the sum of the norms. In principle, the functional norm can be anything but in this case we used the 𝓁1 norm (the absolute area under the curve),

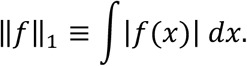

The for an n-member ensemble, the consensus is then

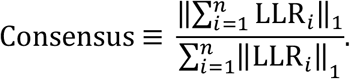

For the *unsigned consensus*, we first generate every permutation of the LLRs used and their inverses, -LLR, up to an overall sign. For example, for a pair of LLRs there are only two options,

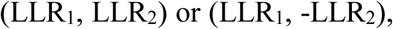

and for three LLRs there are four options,

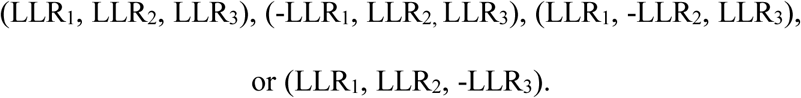

The consensus is then calculated over each these sets and the maximum value is taken to be the value of the unsigned consensus.

### Recurrent neural network model

#### 1. Network elements

We construct a network of N recurrently connected “firing rate” model neurons to perform a facsimile of the auditory discrimination task. Each model neuron is characterized by an activation variable *x*_*i*_ and nonlinear response function *φ*(*x*) = tanh (*x*) that describes its firing rate. *N* = 1000 neurons for **Fig. 7**. Our results are independent of network size (tested on a range from N=1000-10,000) and instead depend on the relative fraction of neurons with specific response properties (to be described).

We denote the connections between model neurons by the *NxN* synaptic weight matrix 𝑱 (**Fig. 7a**) The individual synaptic weights are initially chosen independently and randomly from a Gaussian distribution with mean and variance given by ⟨*𝑱*_*ij*_⟩ = 0 and 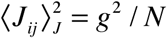, where g sets the initial strength of recurrent synapses. We used *g* values between 0.5 and 0.7 so that units are weakly connected prior to training. The entries of ***J*** are either held fixed or modified during training depending on the fraction of plastic synapses^41^.

The activation variable for each network neuron x¿ is determined by

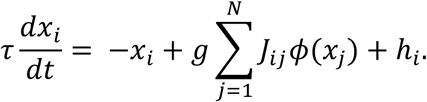

In the above equation, τ = 10 ms is the time constant of each unit in the network which sets the time scale of network dynamics and *h*_*i*_ is the external input to unit *i*. The network equations are integrated using Euler method with an integration time step, *dt* = 1 ms.

#### 2. Design of Inputs and Network Outputs

Auditory input, 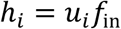 is provided to 10% of the network units through the vector of input weights, **u**. To represent the auditory stimuli used in the experiment, *f*_in_ is modeled as a square pulse of duration 100 ms and amplitude proportional to the frequency of the sound used experimentally, ranging from 0.5 kHz to 32 kHz in octave increments (see left panel of **Fig. 7a**). On each trial the network receives one of these “auditory” inputs. After network training, it should respond only to the “target tone” of 4 kHz with an output pulse modeled by *f*_out_(*t*) and produce no response to the other, “non-target tones” (see right panel of **Fig. 7a**).

The output of the network *z*(*t*) is defined as a sum of unit firing rates from the 90% of the units not directly driven by the tone, and weighted by a vector of readout weights, **w**, such that 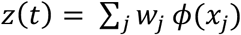 Successful performance is achieved by matching *z*(*t*) to the overall output *f*_out_(*t*) by modifying the readout weights **w** using recursive least squares^38^ (see also section 3 below).

In addition to the auditory inputs *h*_*i*_, a fraction (75% of *N)* of the network units sometimes receive stochastic noise during training to ascertain through decoding that they were nominally non-responsive units (see section 3 below). This injected noise varies randomly and independently at every time step as a Gaussian random variable between 0 and 1, drawn from a zero mean and unit variance distribution. Its amplitude is scaled by 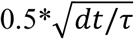.

#### 3. Network Training and Evaluating Performance

During training, inputs of individual neurons whose weights are plastic are compared directly with target functions to compute a set of error functions. In fullFORCE^38^, these target functions are generated by the units in an almost identical, auxillary network in which there are no plastic synapses. However, this network receives the target output *f*_out_(*t*) as an external input, which allows it to perform the task trivially. The recurrent connections of the auxillary network are also sampled as described above, but with a *g* value between 1.2 and 1.5, which have been shown to be effective for this method^41^.

Only the weights between 25% of the network units are modified through fullFORCE^41^ (typically picked at random from the network, but in the cases when noise is used during training, these are noise-free). During training, these synapses undergo modification by a recursive least squares procedure, comparing the activity of the selected units to the activity of neurons in the auxiliary network. As a result of this partial reconfiguration of the network, the trained units evolve into the subpopulation of responsive (R) units, whereas the remaining 75% remain nominally non-responsive (NNR). Units are confirmed to be nominally non-responsive using the following two methods: first, in noise-free simulations, by verifying that their activity fluctuations remained below a threshold of 10% (responsive neurons demonstrated fluctuations greater than this threshold); and second, by attempting to decode the output from the activity of different subpopulations.

The convergence of the fullFORCE algorithm was assayed by 1) directly comparing the output of the network *z*(*t*) with the target function, *f*_out_(*t*), and 2) by calculating and tracking the squared error between the individual unit rates and their targets, both during training and at the end of the simulation. Starting from a random initial state, each network is trained for 500 trials of duration 300 ms (3,000 time steps) in **Fig. 7**.

The behavioral performance of the trained network is computed as a percentage of trials in which the network produced a pulsed output (green trace in the right panel of **Fig. 7a**) in response to the target tone (input schematized in the left panel of **Fig. 7a**) and produced 0 in response to the non-target tone (purple trace in right panel of **Fig. 7a**). A typical trained network of 1000 neurons was able to perform the task to a percent correct level of 100% (**Fig. 7b**). Performance was evaluated in the same way after silencing neurons in the responsive or nominally nonresponsive cohort, or alternatively, in a randomly selected fraction of network neurons (**Fig. 7c,d**).

The predicted decrease in decoding performance for a random subset of the network seen in panel **Fig. 7d** (*D*_pred_.) was derived from the psychometric inactivation curves of each subpopulation treated independently. *D*_pred_. was calculated by assuming the decrease in decoding performance is a linear sum of the decreases due to the inactivated responsive cells (*D*_R_) and the decreases due to the nonresponsive cells (*D*_NNR_) present in the mixed ensembles,

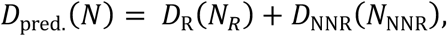

where *N* is the total number of neurons inactivated and *N*_R_ (*N*_*NNR*_) are the expected number of responsive (nominally non-responsive) units included in the randomly inactivated fraction (In our case, *N*_*R*_ = 0.25 *N* and *N*_*NNR*_ = 0.75 N).

### Analysis of selection rule encoding from Rodgers & DeWeese 2014

Using our novel ISI-based decoding algorithm, we analyzed cells found to be non-responsive in a previously published study^5^. Briefly, rats were trained on a novel auditory stimulus selection task where animals had to respond to one of two cues while ignoring the other depending on the context. Rats held their nose in a center port for 250 to 350 ms and were then presented with two simultaneous sounds (a white noise burst played from only the left or right speaker and a high or low pitched warble played from both speakers). In the “localization” context animals were trained to ignore the warble and respond to the location of the white noise burst and in the “pitch” context they were trained to ignore the location of the white noise burst and respond to the pitch of the warble. Cells recorded from both primary auditory cortex and prefrontal cortex (prelimbic region) were shown to be responsive to the selection rule during the pre-stimulus period (i.e. firing rates differed between the two contexts). Non-responsive cells were reported but not further analyzed.

We confirmed that these nominally non-responsive cells satisfied our own positive statistical criteria for non-responsiveness (described above) by comparing their average spiking activity in the 100 ms immediately preceding stimulus onset across contexts. To determine whether the prestimulus activity of nominally non-responsive cells also encoded the selection rule, we decoded the task context (localization vs. pitch) on single-trials using our ISI-based decoding algorithm. Cells shown in **Fig. 5b** were deemed statistically significant when compared to the decoding performance of a control using synthetically generated data (p<0.05).

### Statistical analysis

All statistical analyses were performed in Python, MATLAB, or GraphPad Prism 6. Datasets were tested for normality, and appropriate statistical tests applied as described in the text (e.g., Student’s paired t-test for normally distributed data, Mann-Whitney U test and Wilcoxon matched-pairs signed rank test for non-parametric data).

## Acknowledgements

We thank E. Simoncelli, N.D. Daw, C.S. Peskin, A.A. Fenton, E. Kelemen, K. Kuchibhotla, E. Morina, M. Aoi, and A. Charles for comments, discussions, and technical assistance, and C.A. Loomis and the NYU School of Medicine Histology Core for assistance with anatomical studies. Shari E. Ross produced the illustration in **Fig. 1a**. This work was funded by an NYU Provost’s Postdoctoral Fellowship and NIDCD (DC015543-01A1) to M.N.I.; a NARSAD Young Investigators Award, NIMH (T32), and NIMH (KMH106744A) to I.C.; a James McDonnell Understanding Human Cognition Scholar Award to K.R; a Fordham University Grant for Cloud Based Computing Research Projects to B.F.A.; and NIDCD (DC009635 and DC012557), the NYU Grand Challenge Award, a Howard Hughes Medical Institute Faculty Scholarship, a Hirschl/Weill-Caulier Career Award, and a Sloan Research Fellowship to R.C.F. The authors declare no competing financial interests.

## Author Contributions

M.N.I., I.C., and R.E.F. collected the data. M.N.I and B.F.A. analyzed the data, M.R.D. and B.D. verified the analysis, K.R. performed network modeling, and M.R.D. and C.C.R. provided additional datasets. M.N.I., B.F.A., R.C.F. designed the study. M.N.I, B.F.A. and R.C.F. wrote the paper.

## Author Information

The authors declare no competing financial interests. Correspondence and requests for materials should be addressed to.

## Supplementary Figures

**Supplementary 1.**
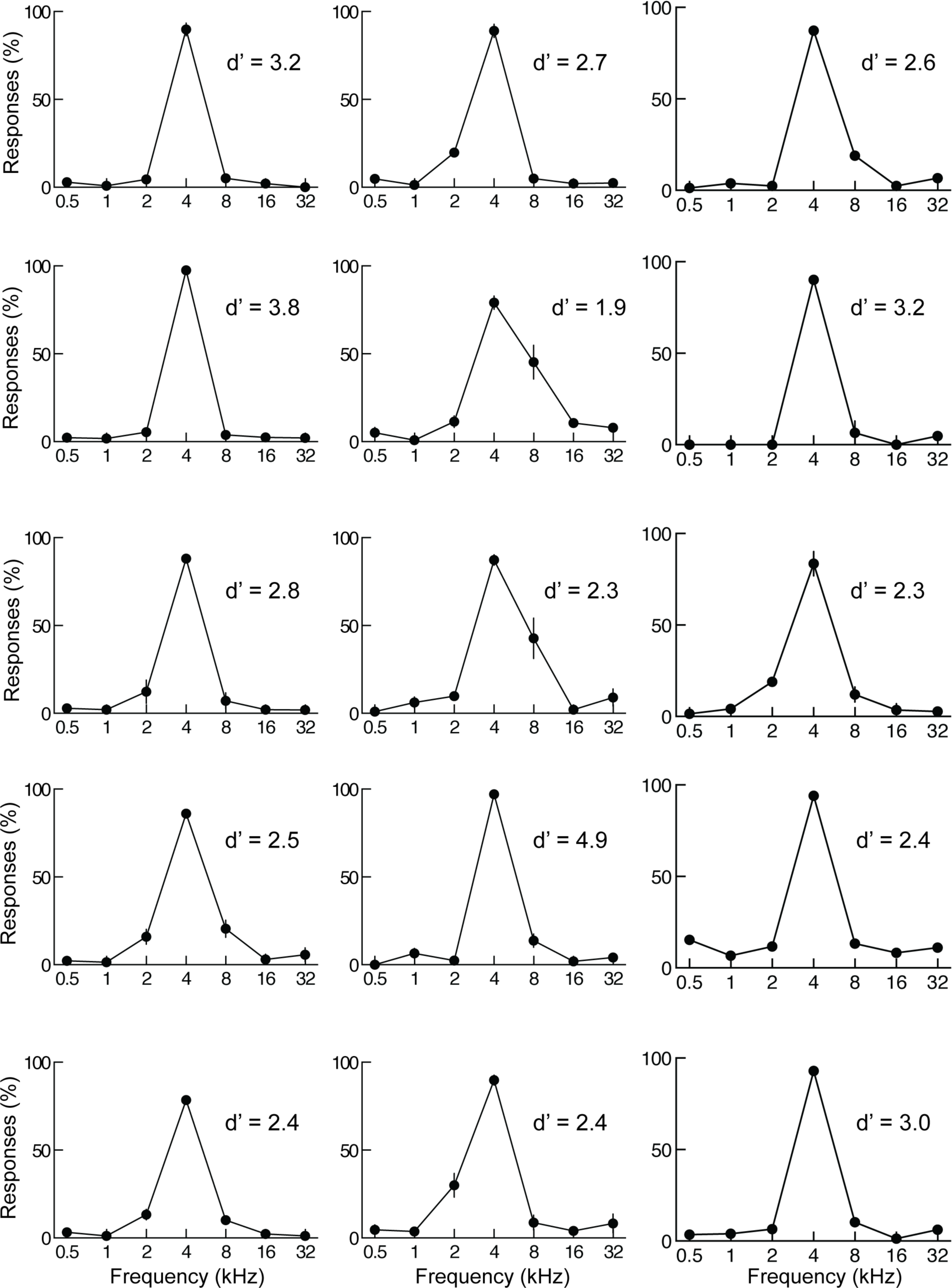
Individual response curves from 15 animals included in this study. Each panel shows data from a different animal including behavioral d’ for distinguishing target from non-target tones. We used a criteria of d’ ≥1 for inclusion in this study. Response curves here are for an average of 3-4 sessions. Error bars represent S.E.M.

**Supplementary 2.**
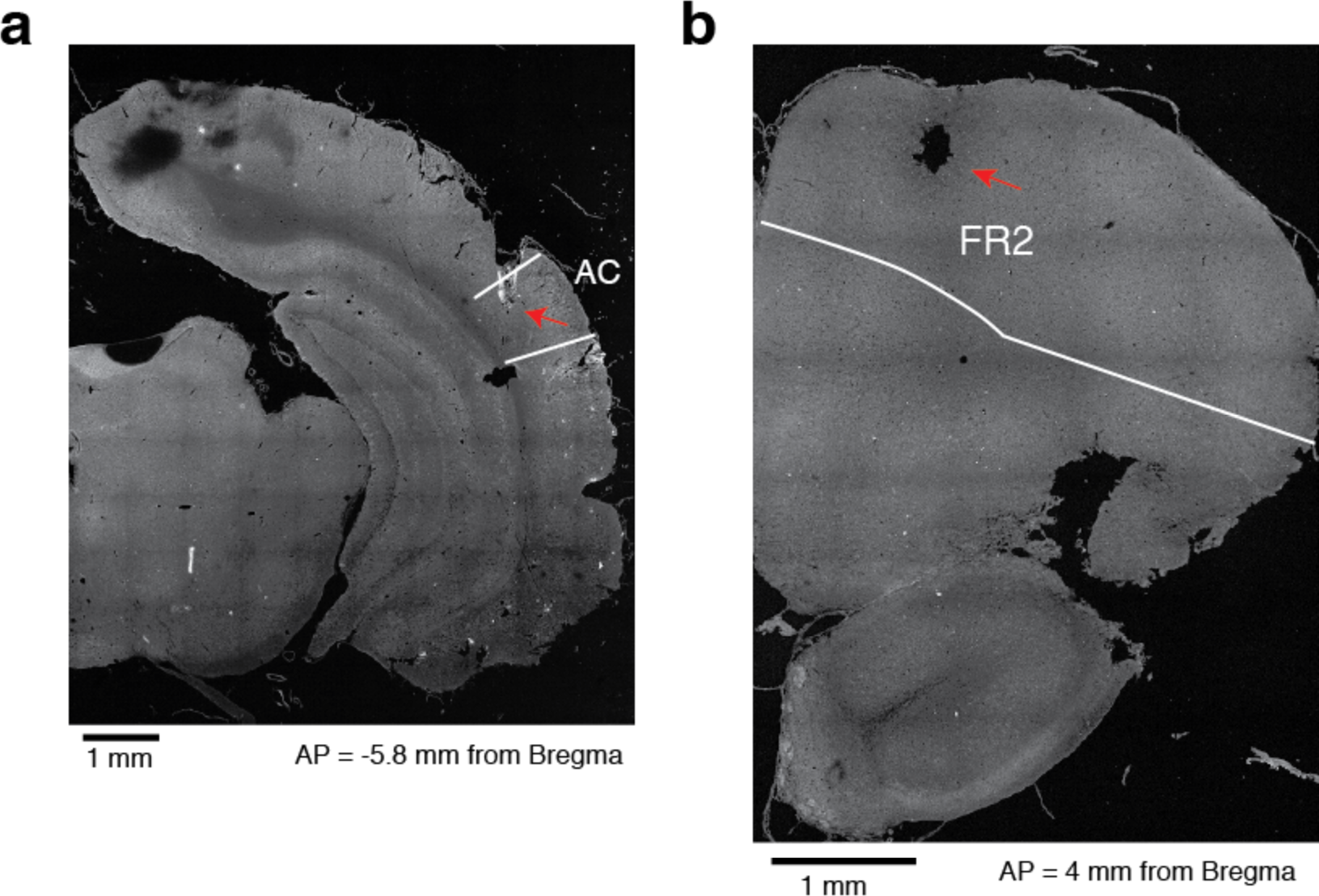
Histological placement of cannulas in AC and FR2. **a**. Example of a coronal section of a rat implanted with cannulas in primary auditory cortex (AC). The white lines represent the borders of AC^38^. **b**. Example of a coronal section of a rat implanted with cannulas in FR2. The white lines represent the borders of FR2^38^.

**Supplementary 3.**
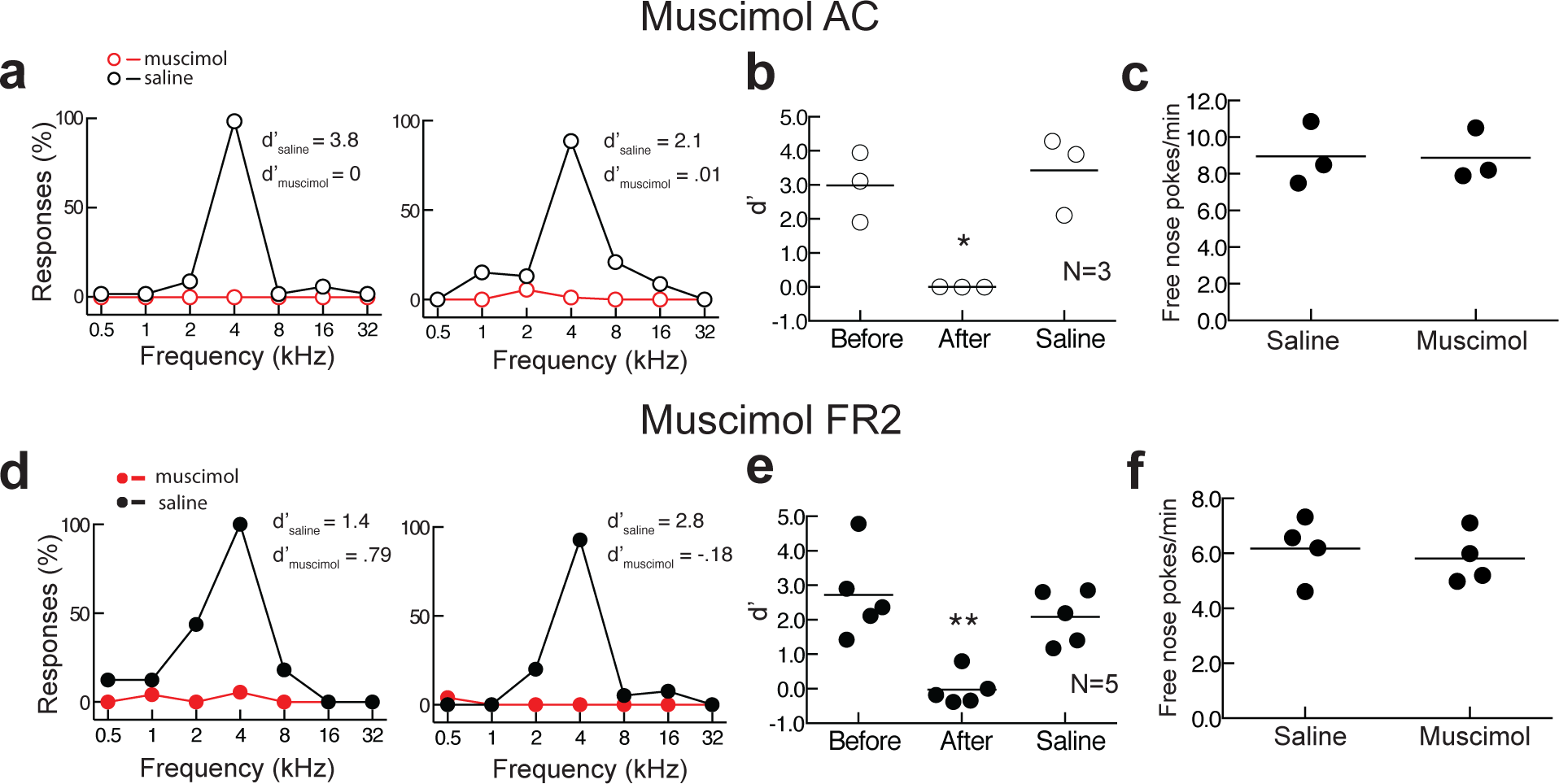
Bilateral infusion of muscimol into either AC or FR2 significantly impairs task performance. **a**. Behavioral performance after muscimol infusion (red) or saline control (black) in AC from two individual animals. **b**. Summary of performance on day before infusion, after muscimol infusion into AC, and after saline control infusion (N=3 animals). Performance was impaired after muscimol infusion (p=0.03 Student’s paired two-tailed t-test, *p <0.05). **c**. Behavior of one animal allowed to freely nose poke for food without tones being presented. This behavior was not affected by muscimol inactivation (average of 3 sessions, p>0.99 Wilcoxon matched-pairs signed rank test). Error bars represent S.E.M. **d**. Behavioral performance for two animals infused bilaterally with muscimol into FR2. **e**. Summary of performance before, during, and after muscimol infusion into FR2 (N=5 animals). Performance was impaired after muscimol infusion (p=0.009 Student’s paired two-tailed t-test, **p<0.01). **f**. Muscimol in FR2 did not impair free nose poking for food without tones being presented in two animals (average of 4 sessions, p=0.62, Wilcoxon matched-pairs signed rank test).

**Supplementary 4.**
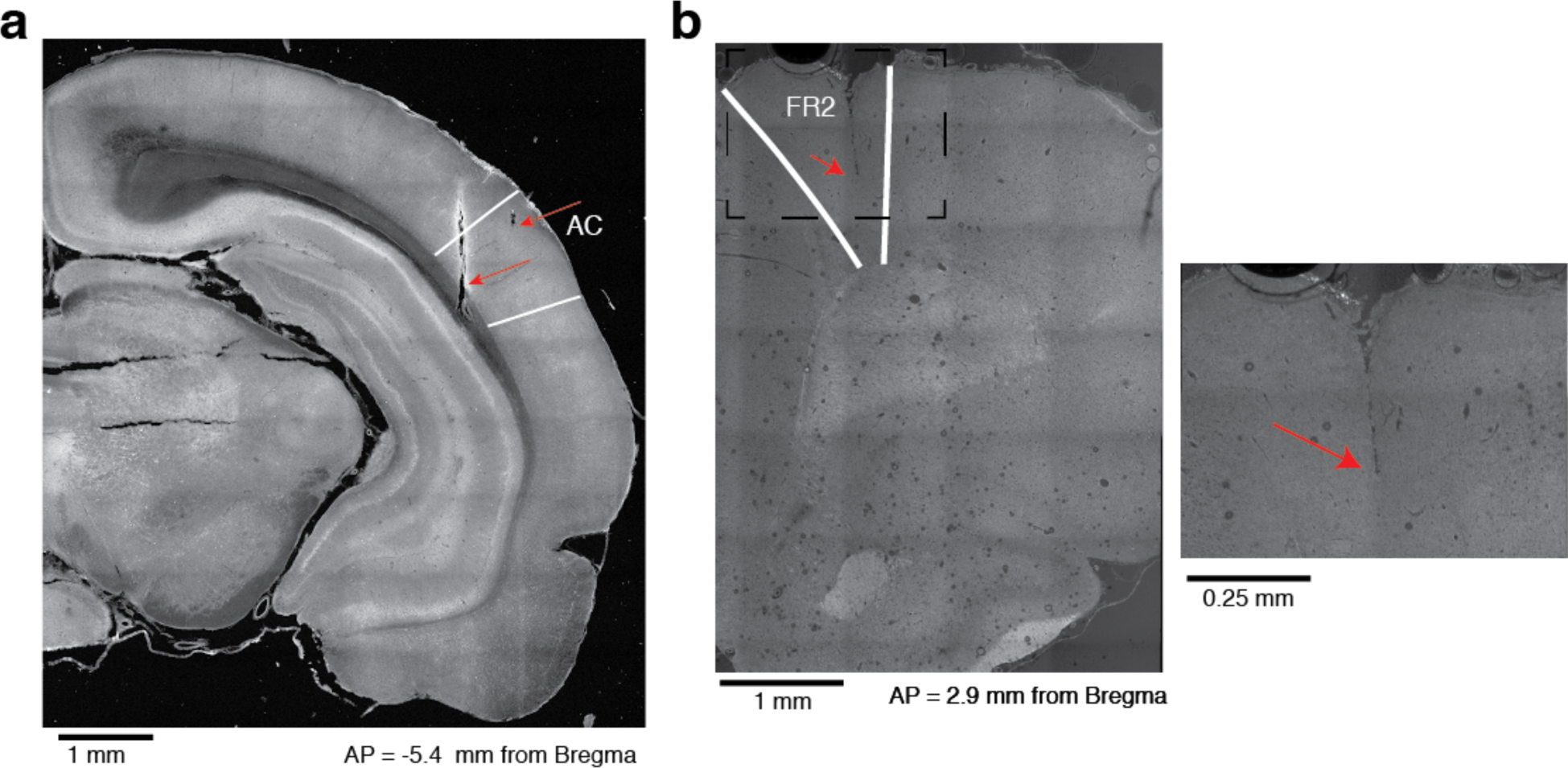
Histological placement of electrodes in AC and FR2. **a**. Example of electrode tracks and electrolytic lesions in AC. The white lines represent the borders of AC. **b**. Example of an electrode track in FR2. The white lines represent the borders of FR2. Left, section imaged at 10X. Right, the same section imaged at 40X.

**Supplementary 5.**
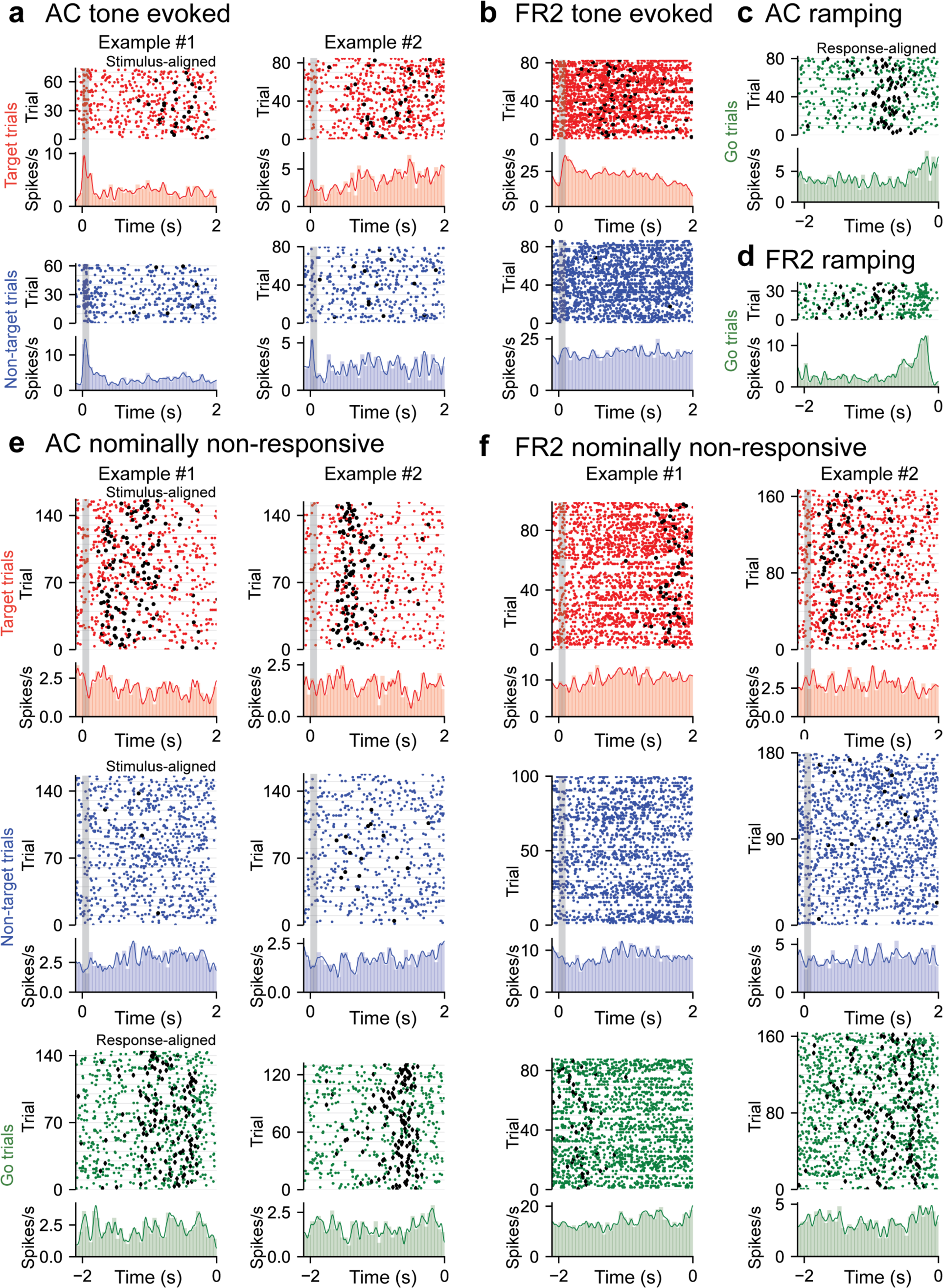
Examples of tone evoked, ramping, and nominally non-responsive cells from AC and FR2. **a**. Two example tone-evoked cells recorded from AC. Rasters and PSTHs of target (red) and non-target (blue) trials shown. Stimulus shown as grey bar and black circles represent behavioral response. (Example #1: average evoked spikes on target tones = 0.55, on nontarget = 0.92. Example #2: average evoked spikes on target tones = 0.096, on non-target = 0.12; note that example #2 is only non-target tone evoked). **b**. Example target tone-evoked cell recorded from FR2. Rasters and PSTHs of target (red) and non-target (blue) trials shown (average evoked spikes on target tones = 0.37, on non-target = 0.20). **c**. Example ramping cell recorded from AC. Rasters and PSTH of go trials (green) shown (ramp index = 2.8). **d**. Example ramping cell recorded from FR2 (ramp index = 4.9). Rasters and PSTH of go trials (green) shown. **e**. Two example nominally non-responsive cells recorded from AC. Rasters and PSTH of target (red), non-target (blue), and go (green) trials shown. (Example #1: average evoked spikes on target tones = 0.12, on non-target = −0.12, ramp index = −0.85; p_tone_<0.001, p_ramp_=0.010, 2,000 bootstraps; Example #2: average evoked spikes on target tones = −0.020, on non-target = −0.038, ramp index = 1.1; ptone<0.001, pramp=0.004, 2,000 bootstraps). f. Two example nominally non-responsive cells recorded from FR2 (Example #1: average evoked spikes on target tones = 0.081, on non-target = −0.15, ramp index = 1.4; p_tone_<0.001, p_ramp_=0.019, 2,000 bootstraps; Example #2: average evoked spikes on target tones = 0.046, on non-target = −0.070, ramp index = 1.8; p_tone_<0.001, p_ramp_=0.033, 2,000 bootstraps). Rasters and PSTH of target (red), non-target (blue), and go (green) trials shown.

**Supplementary 6.**
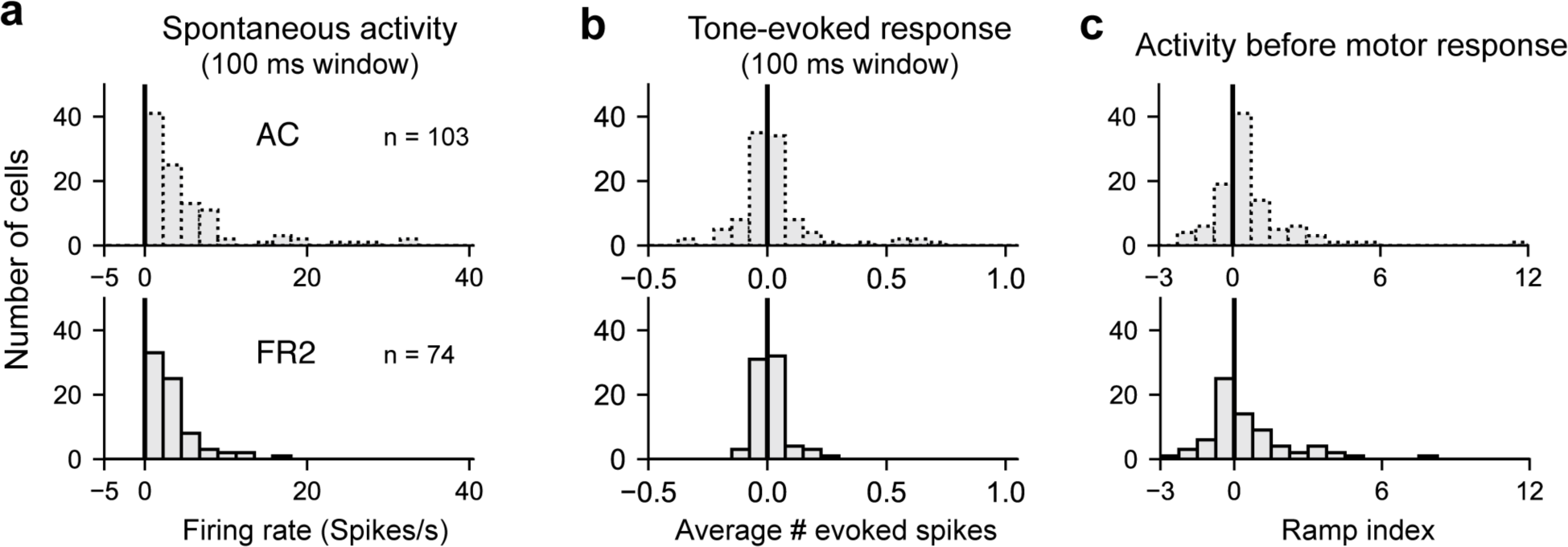
Histograms of **a**. spontaneous firing rate, **b**. average number of tone-evoked spikes, and c. ramp index for AC (top) and FR2 (bottom).

**Supplementary 7.**
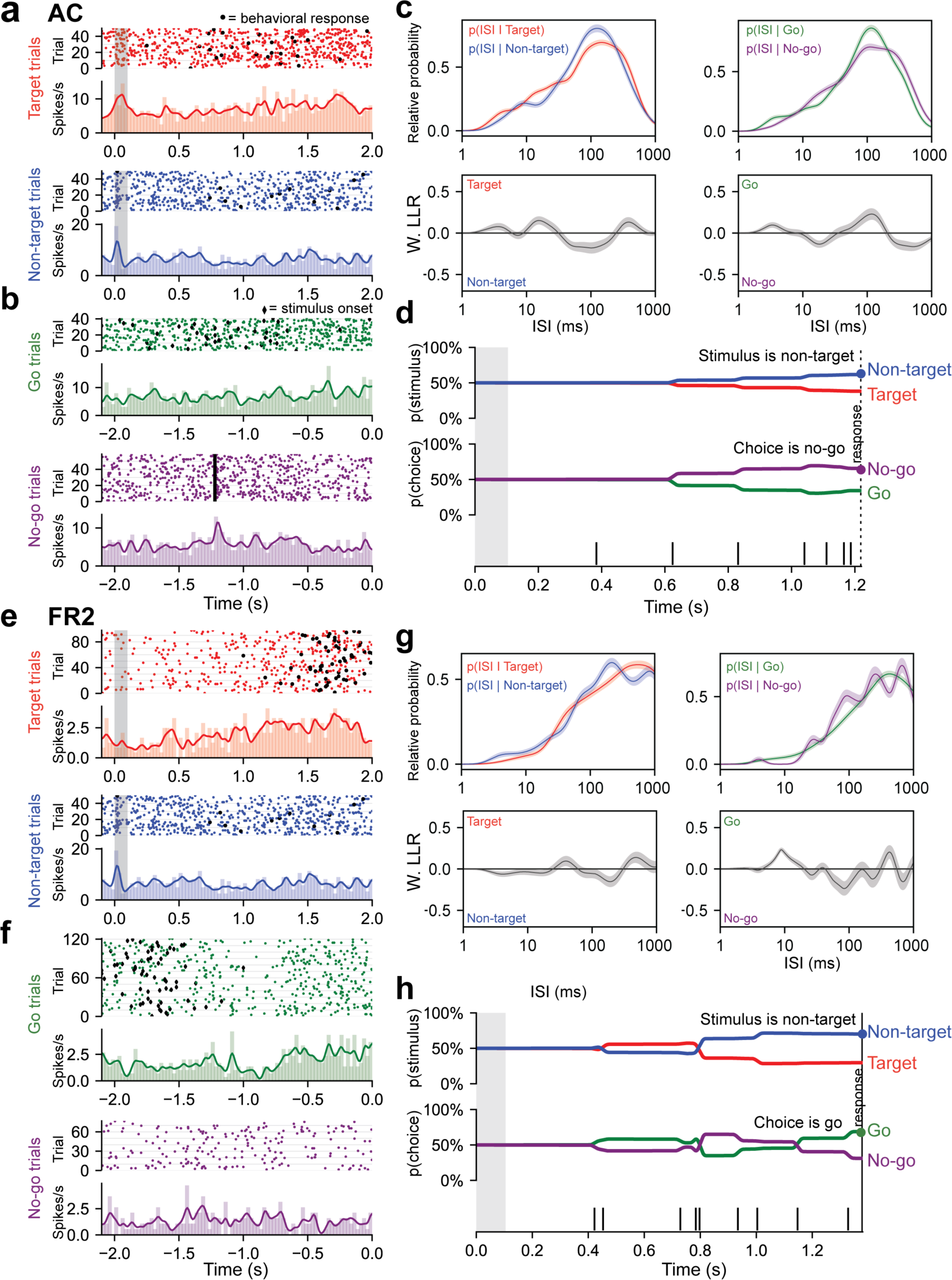
Decoding algorithm to determine stimulus category and choice in single-unit ISIs from AC and FR2 for two additional neurons. **a-d**. Decoding algorithm applied to a sample neuron in AC. **a**. Single-unit activity sorted by stimulus condition: target trials (red) and non-target trials (blue). Black circles represent the behavioral response. **b**. Trials aligned to behavioral response: go (green) and no-go (purple). Black diamonds in both go and no-go trials represent stimulus onset. **c**. All ISIs during the trial (following stimulus onset and before behavioral choice) are aggregated into libraries for each condition (average response time is used on no-go trials). Probability of observing a given ISI on each condition was generated by using Kernel Density Estimation on the libraries from **a**. Top left are target (red) and non-target (blue) probabilities and on right are go (green) and no-go (purple). Below left (right) are the log likelihood ratios (LLR) for the ISIs conditioned on stimulus category (behavioral choice). When curve is above zero the ISI suggests target (go); when it is below zero the ISI suggests non-target (no-go). **d**. Probability functions from c. were used as the likelihood function to estimate the prediction of a spike train on an individual trial (bottom). Bayes’ rule was used to update the probability of a stimulus (top) or choice (bottom) as the trial progresses and more ISIs were observed. Prediction for the trial was assessed at the end of the trial as depicted by the highlighted dot. **e-h**. as in **a-d** except the decoding algorithm is applied to a neuron from FR2.

**Supplementary 8.**
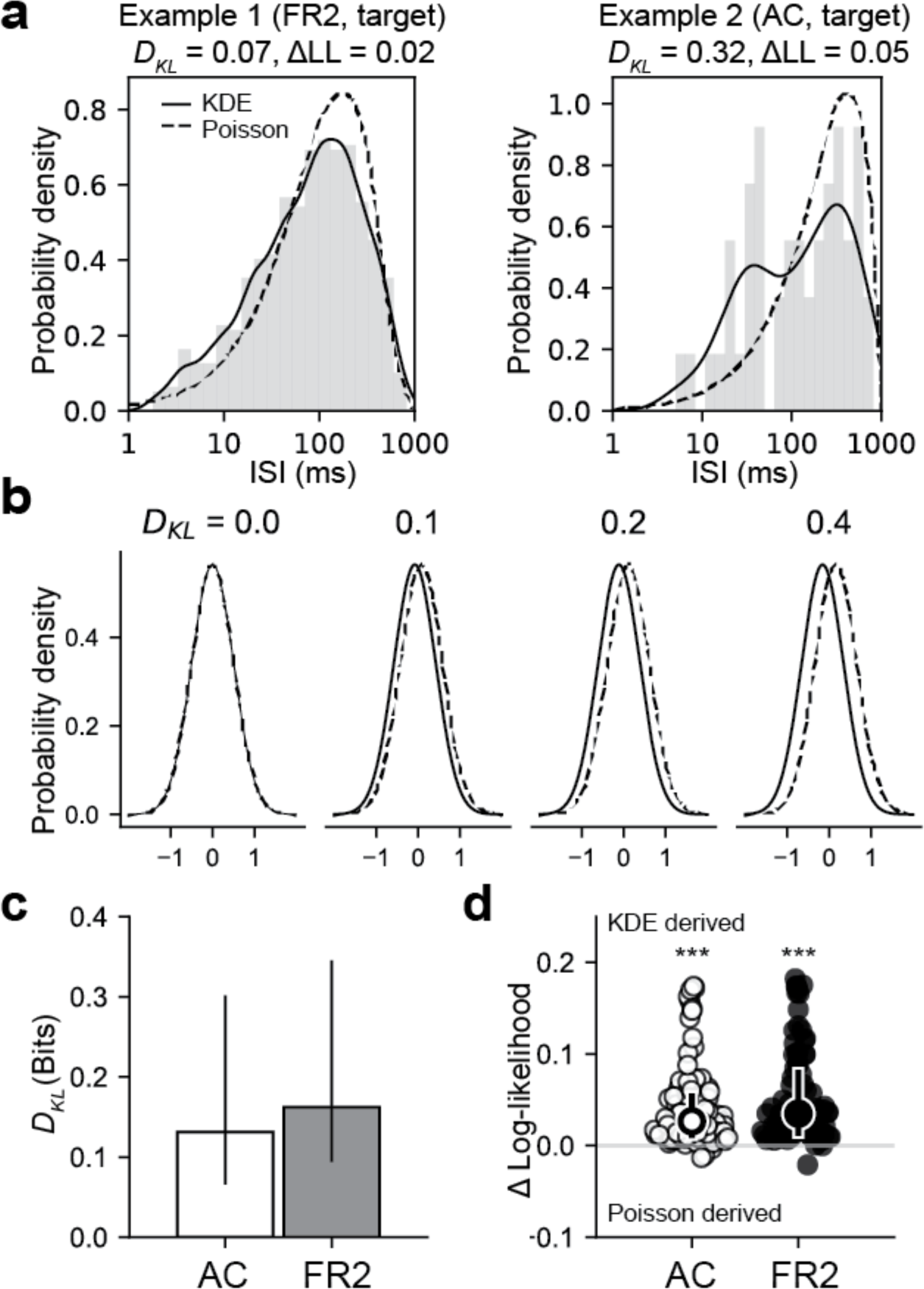
Empirical ISI distributions are better modeled using non-parametric methods. **a**. ISI histograms from two example cells on target trials with the corresponding non-parametric Kernel Density Estimate (KDE) distribution (solid lines) and the distribution derived from a rate-modulated Poisson process (dashed lines). Above each example is the Kullback-Leibler divergence (DKL) quantifying the difference between these two distributions, and the difference in the average log-likelihood of the data (ΔLL) where positive values indicate that the data is better described by the non-parametric KDE distribution. **b**. Constructed examples of the KL divergence for four pairs of normal distributions with equal standard deviations and various mean offsets as a visual reference **c**. Summary of all KL divergence values for both stimulus and choice in AC (white) and FR2 (grey). Bar indicates median and error bars indicate bottom and top quartiles. **d**. Summary of difference between log-likelihood of observed data under non-parametric KDE and rate-modulated Poisson distributions. Positive values indicate KDE distributions are generically a superior fit for the data (AC: p = 1.1×10^−15^ FR2: p = 1.2×10^−16^, Wilcoxon signed-rank test).

**Supplementary 9.**
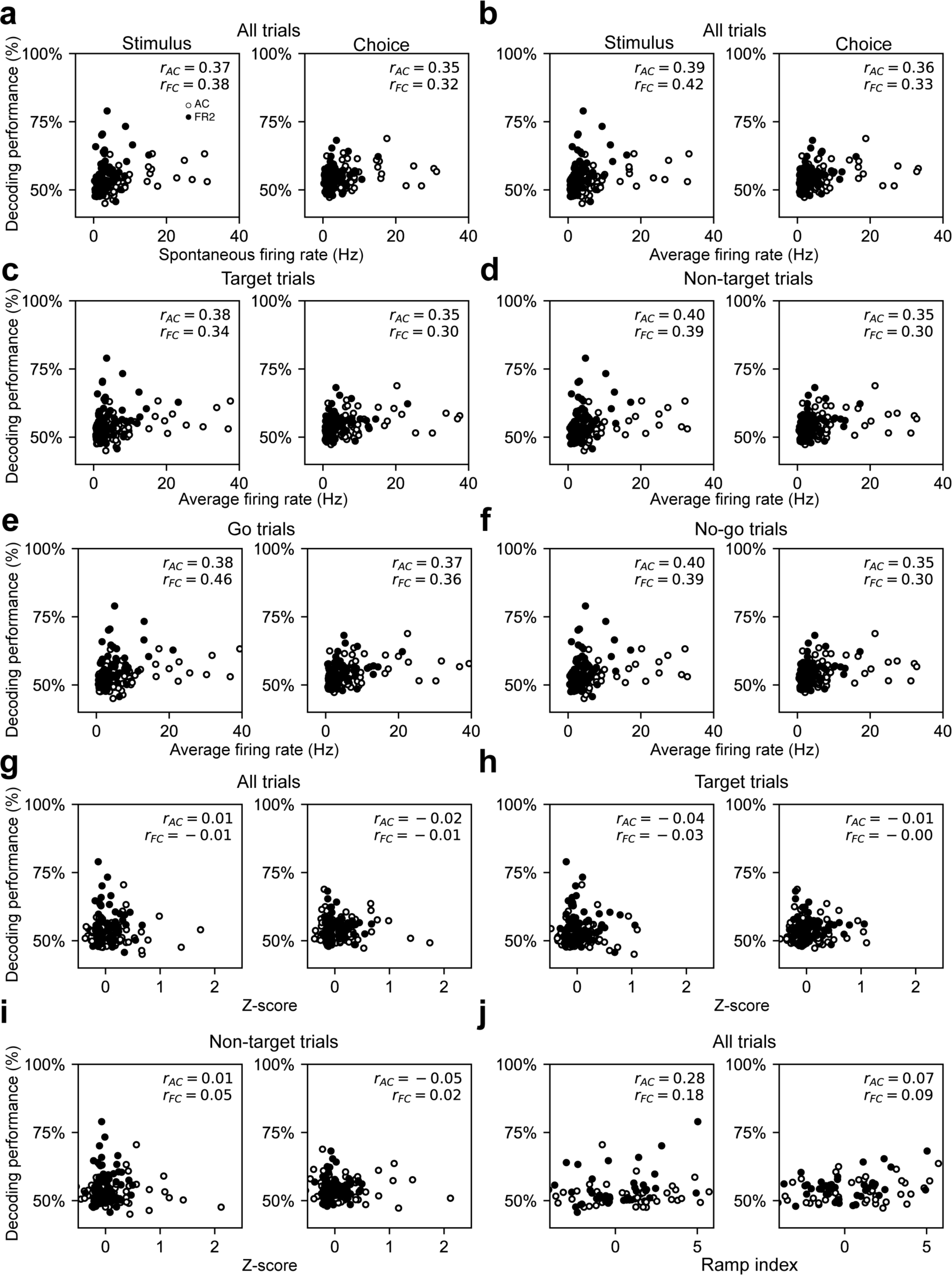
Lack of correlations between classical firing rate metrics and stimulus or choice decoding performance. **a**. Stimulus and choice decoding performance versus the spontaneous firing rate for both target and non-target trials, (r_ac_ = 0.37, 0.35; r_FR2_ = 0.38, 0.30). **b**. Stimulus and choice decoding performance versus average firing rate for both target and non-target trials, (r_ac_ = 0.39, 0.36; r_fr2_ = 0.42, 0.33). **c**. Stimulus and choice decoding performance versus average firing rate for target trials only, (r_ac_ = 0.38, 0.35; r_fr2_ = 0.34, 0.30). **d**. Stimulus and choice decoding performance versus average firing rate for non-target trials only, (r_ac_ = 0.40, 0.35; r_fr2_ = 0.39, 0.30). **e**. Stimulus and choice decoding performance versus average firing rate for go trials only, (r_ac_ = 0.38, 0.37; r_fr2_ = 0.46, 0.36). f. Stimulus and choice decoding performance versus average firing rate for no-go trials only, (r_ac_ = 0.40, 0.35; r_fr2_ = 0.39, 0.30). **g**. Stimulus and choice decoding performance versus z-score for all trials, (rAC = 0.01, −0.02; rFR2 = 0.01, −0.01). h. Stimulus and choice decoding performance versus z-score for target trials only, (r_ac_ = −0.04, −0.01; r_fr2_ = −0.03, −0.002). **i.** Stimulus and choice decoding performance versus z-score for non-target trials only, (r_ac_ = 0.01, −0.05; r_fr2_ = 0.05, 0.02). **j.** Stimulus and choice decoding performance versus ramp index, (r_ac_ = 0.28, 0.07; r_fr2_ = 0.18, 0.09).

**Supplementary 10.**
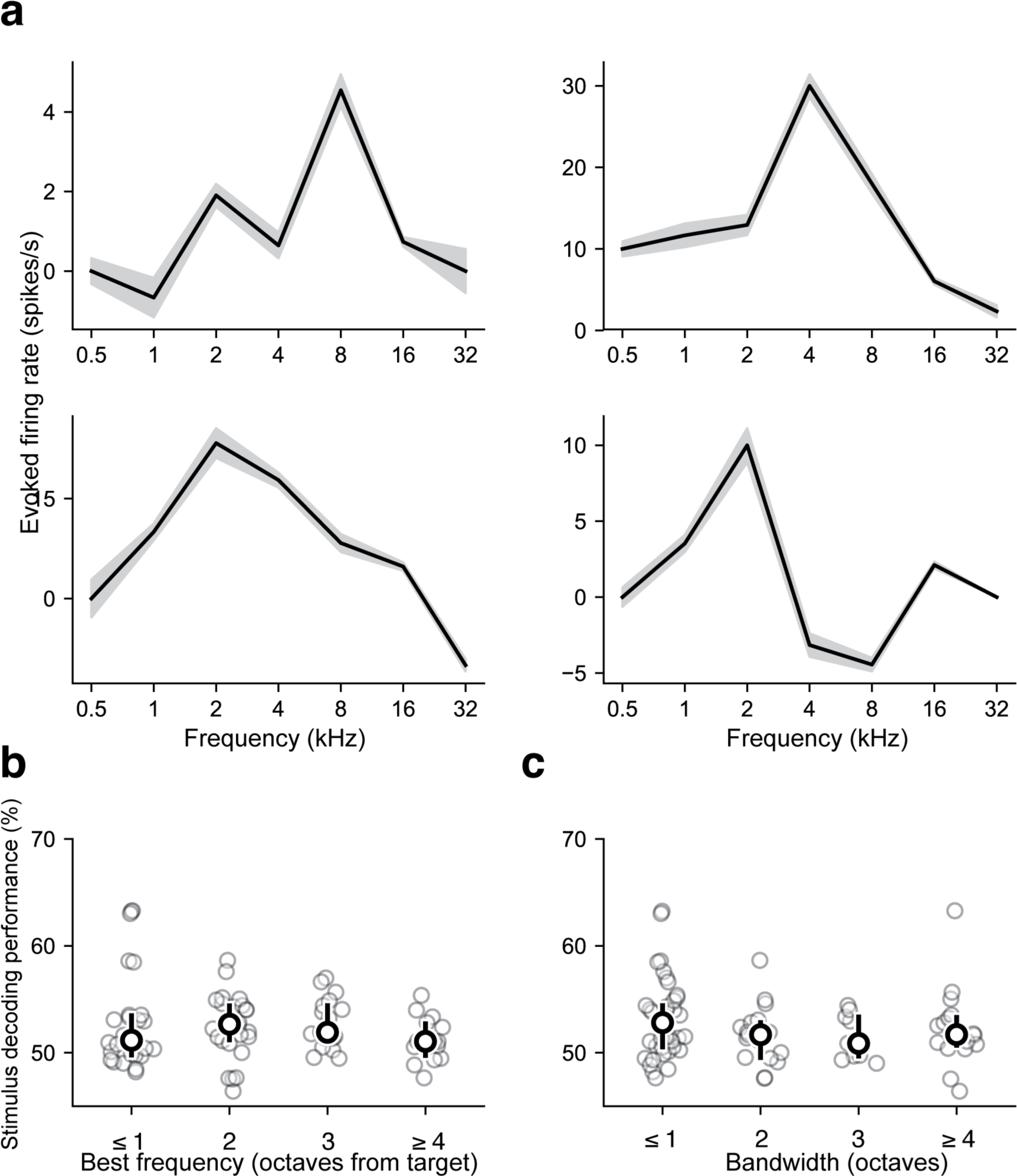
Stimulus decoding in AC independent of receptive field properties. **a**. Examples of tuning curves from four different neurons constructed from responses in AC. Gray regions represent S.E.M. **b**. Stimulus decoding performance as a function of best frequency as measured relative to the target tone frequency. No significant differences were found between groups (p>0.2, Mann Whitney U test, two-sided). **c**. Stimulus decoding performance as a function of receptive field bandwidth tuning. No significant differences were found between groups (p>0.1, Mann Whitney U test, two-sided).

**Supplementary 11.**
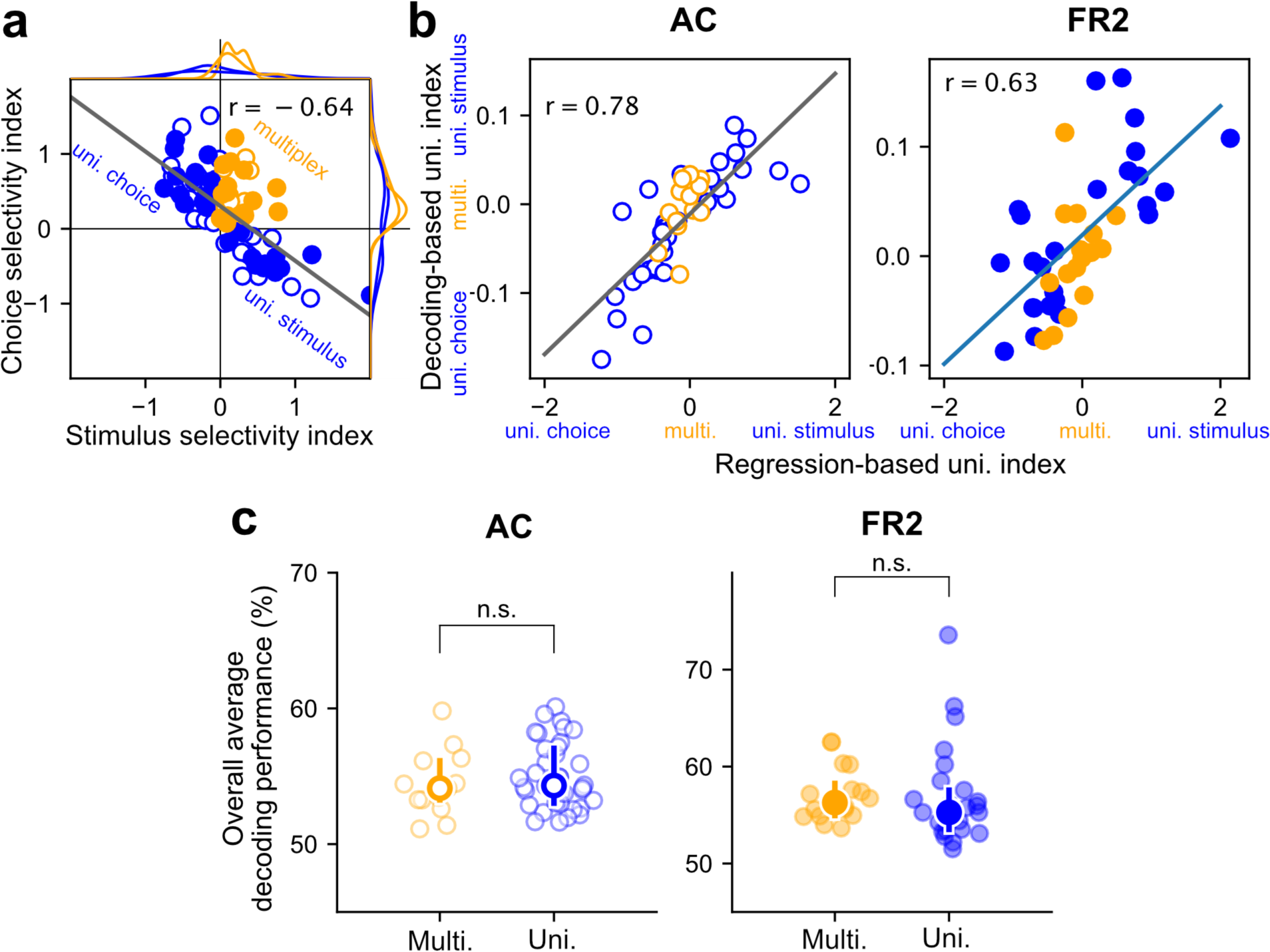
Decoding performance is a sufficient measure of uni/multiplexing. Given the correlation between stimulus category and behavioral choice we used a regression based analysis to determine whether decoding performance alone was sufficient to establish whether cells were multiplexed for both behavioral variables. We used multiple regression to create an alternative definition of multiplexing and uniplexing and then demonstrated this definition coincides with the one used in the paper based solely on decoding performance. **a**. Choice selectivity index versus stimulus selectivity index for single cells. Each index quantifies the extent to which the corresponding variable was predictive of decoding performance. Multiplexed cells (orange symbols) have positive values on both indices. Uniplexed cells (blue symbols) are only positive for one of the two indices. Each cell was projected on the linear regression (grey line) to construct a regression-based uniplexing index. Multiplexed cells were close to zero on this measure and cells uniplexed for stimulus or choice were positive or negative respectively. **b**. The decoding-based uniplexing index (difference between stimulus and choice decoding performance) versus the regression-based index defined in a for AC (left, open symbols) and FR2 (right, filled symbols). In both regions, these two measures of uni/multiplexing were correlated. c. Overall decoding performance (average of stimulus and choice decoding) for multiplexed cells versus uniplexed cells in AC (left) and FR2 (right). There were no systematic differences in decoding performance between multiplexed and uniplexed units (n.s. p_ac_=0.22, pFR2=0.11, Mann-Whitney U test, two-sided).

**Supplementary 12.**
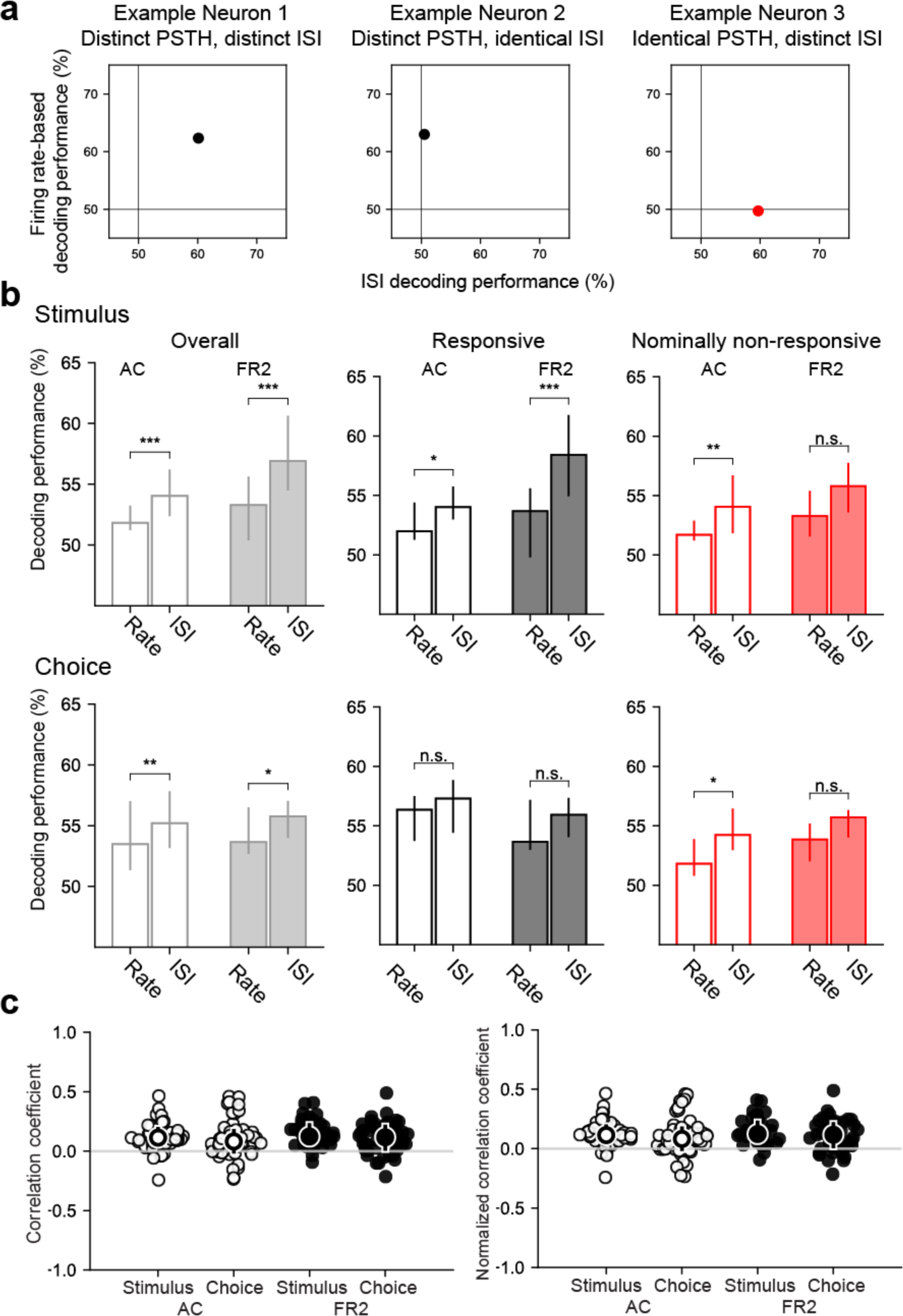
Information captured by ISI-based decoder distinct from conventional rate-modulated (inhomogeneous) Poisson decoder. **a.** Decoding performance comparison for example neurons shown in Figure *2. Left*, Both the trial-averaged firing rate and the ISI distributions can be used to decode stimulus category for this example neuron. *Middle*, Only the firing rate can be used to decode this example. *Right*, In this case, the ISI distributions can be used to decode even when the trial-averaged firing rate cannot. **b**. Comparison of decoding performance for conventional rate-modulated decoder to our ISI-based decoder. *Top row*, stimulus decoding, *bottom row*, choice decoding. *Left*, Overall comparison for all cells. *Right*, Comparison for responsive and non-responsive cells (Stimulus Overall: ***pAC=0.0001, ***pFR2=8×10^−6^, Stimulus Repsonsive: *pAC=0.031, ***pfr2=4×10^−5^, Stimulus Non-responsive: **pAC=0.0019, n.s. pFR2=0.096, Choice Overall: **pAC=0.0057, *pFR2=0.02, Choice Repsonsive: n.s. pAC=0.031, n.s. pFR2=0.08, Choice Non-responsive: *pAC=0.004, n.s. pFR2=0.19, Wilcoxon signed-rank test). Bar indicates median and error bars designate bottom and top quartiles. **c**. *Left*, Matthews correlation coefficient (MCC) between correct predictions of our ISI-based decoder and a conventional rate-modulated firing rate decoder. A MCC value of 1 indicates each decoder correctly decodes exactly the same set of trials whereas −1 indicates each decoder is correct on complementary trials. Values close to 0 indicate that that the relationship between the decoders is close to chance. Typically, values from −0.5 to 0.5 are considered evidence for weak to no correlation (stimulus median & interquartile range: AC=0.10, 0.09, FR2=0.11, 0.12; choice median & interquartile range: AC=0.06, 0.15, FR2=0.08, 0.17). *Right*, Matthews correlation coefficient (MCC) rescaled by the maximum possible correlation given the decoding performance of each method remains fixed. This control demonstrates that the correlation values are not a result of weak decoding performance for one of the decoding methods (stimulus median & interquartile range: AC=0.11, 0.11, FR2=0.12, 0.15; choice median & interquartile range: AC=0.08, 0.17, FR2=0.11, 0.19).

**Supplementary 13.**
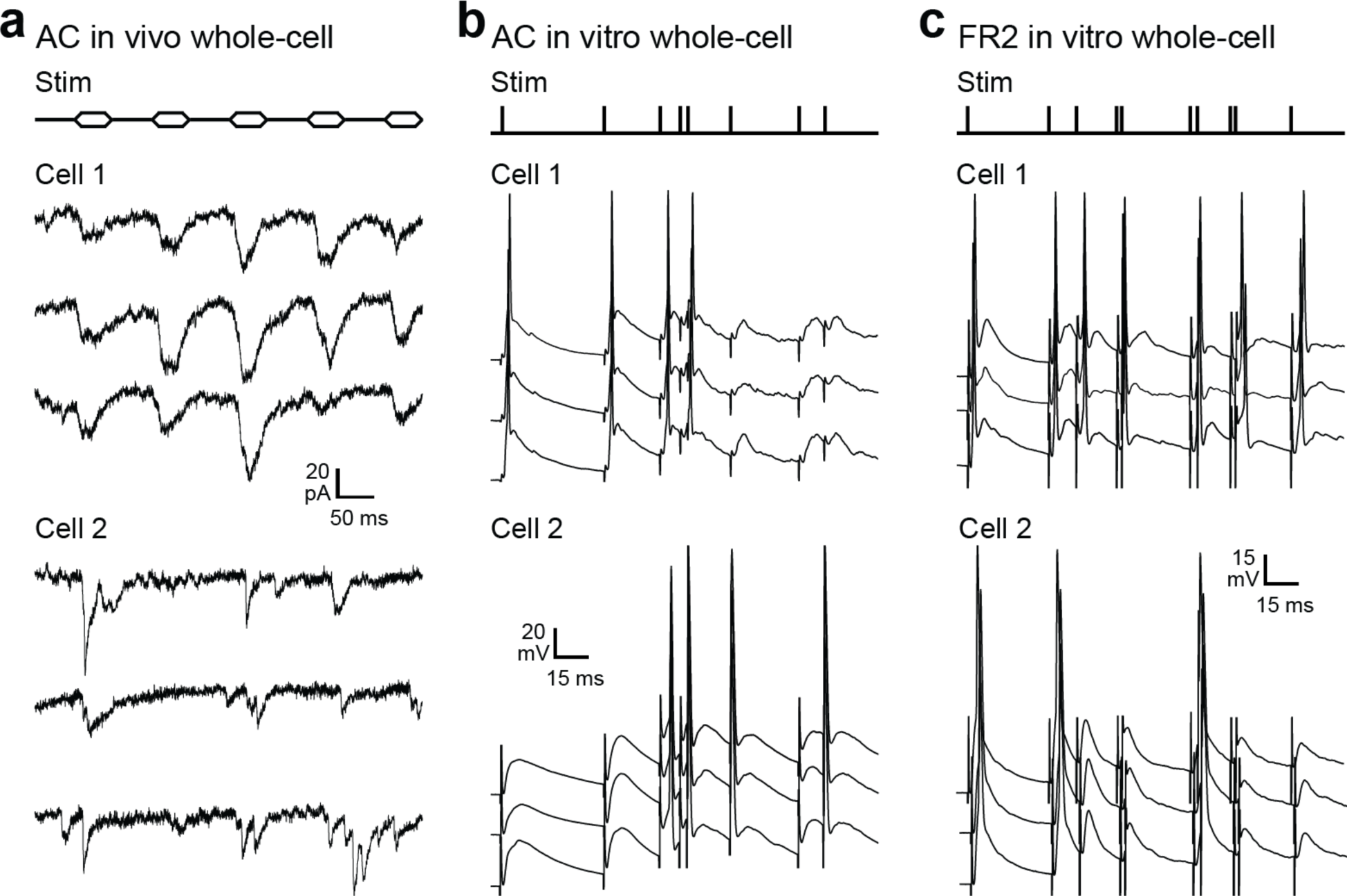
Whole-cell recordings from AC and FR2 neurons showing that different cells can have distinct responses to the same input pattern-necessary for ISI-based decoding by biological networks. In each case, note the reliability of response across trials but differences in response patterns across cells. **a**. Two of eight in vivo whole-cell recordings from anesthetized adult rat primary AC, presenting trains of pure tones at the best frequency for each cell (top, ‘Stim’). **b**. Two of nine whole-cell recordings from adult rat AC in brain slices. Extracellular stimulation was used to present input patterns previously recorded from cortex with tetrode recordings in behaving rats during the auditory task used here, and responses recorded in current-clamp near spike threshold. c. Two of 11 whole-cell recordings from adult rat FR2 in brain slices.

